# Deciphering divergent trypanosomatid nuclear complexes by analyzing interactomic datasets with AlphaFold2 and genetic approaches

**DOI:** 10.1101/2022.11.14.516520

**Authors:** Elvio Rodriguez Araya, Marcelo L. Merli, Pamela Cribb, Vinicius C. de Souza, Esteban Serra

## Abstract

Acetylation signaling pathways in trypanosomatids, a group of early branching organisms, are poorly understood due to highly divergent protein sequences. To overcome this challenge, we used interactomic datasets and AlphaFold2-multimer to predict direct interactions and validated them using yeast two and three-hybrid assays. We focused on MRG domain-containing proteins and their interactions, typically found in histone acetyltransferase/deacetylase complexes. The results identified a structurally conserved complex, *Tc*TINTIN, which is orthologous to human and yeast TINTIN complexes; and another trimeric complex involving an MRG domain, only seen in trypanosomatids. The identification of a key component of *Tc*TINTIN, *Tc*MRGBP, would not have been possible through traditional homology-based methods. We also conducted molecular dynamics simulations, revealing a conformational change that potentially affects its affinity for *Tc*BDF6. The study also revealed a novel way in which an MRG domain participates in simultaneous interactions with two MRG binding proteins binding two different surfaces, a phenomenon not previously reported. Overall, this study demonstrates the potential of using AlphaFold2-processed interactomic datasets to identify protein complexes in deeply branched eukaryotes, which can be challenging to study based on sequence similarity. The findings provide new insights into the acetylation signaling pathways in trypanosomatids, specifically highlighting the importance of MRG domain-containing proteins in forming complexes, which may have important implications for understanding the biology of these organisms and developing new therapeutics. On the other hand, our validation of AlphaFold2 models for the determination of multiprotein complexes illuminates the power of using such artificial intelligence-derived tools in the future development of biology.

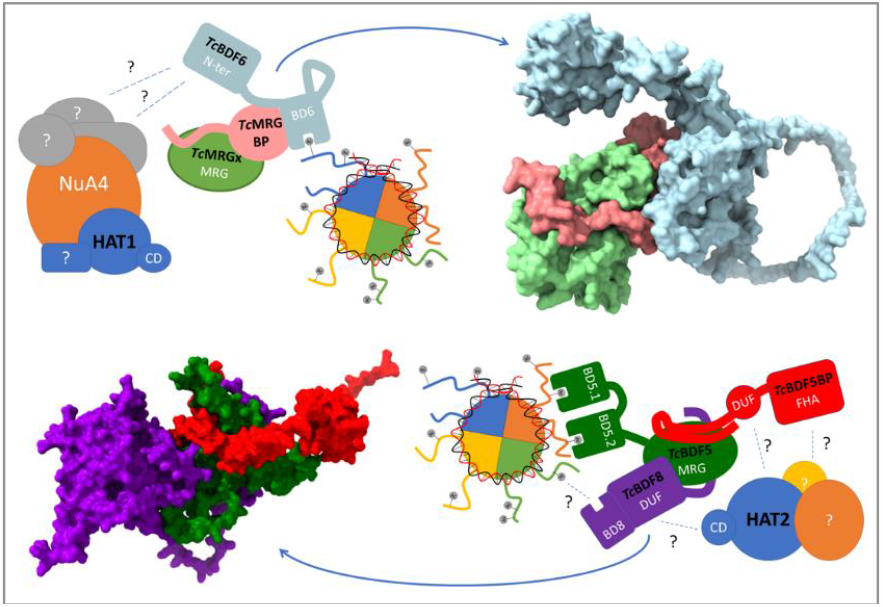

---

Trypanosomatids are protozoans that diverged early in the eukaryotic tree of life, reflected in their highly divergent coding sequences (CDS), sometimes making the search for homology by sequence similarity extremely difficult^1^. Many of them cause human diseases such as sleeping sickness (*T. brucei*)^2^, Chagas disease (*T. cruzi*)^3^, and leishmaniasis (*Leishmania* spp)^4^. They have complex life cycles with different evolutionary stages that range from mammalian hosts to insect vectors. This biological versatility is achieved even with no clear gene transcription regulation mechanisms, given that differential gene expression is not a derivative of gene-specific transcriptional regulation by sequence specific transcription factors^5^. Some promoters have been identified for genes like the SL RNA locus, rRNA locus, procyclin and VSG genes of *T. brucei*^6^. Recently, a promoter sequence was reported in the 5’ region of gene tandem units in *T. brucei*, but the significance of this finding remains yet-to-be well determined^7^. They also contain a set of basal transcription factors that form a very divergent initiation complex^8^. These observations and other evidence have led to hypothesize that transcription regulation in trypanosomatids is mainly epigenetic and gene expression would depend on the degree of chromatin accessibility^9^. Chromatin structure can be altered either by post-translational modifications of the N-terminal tails of histones, by the substitution of one or more histones by their variants, or by the combination of these events. These changes not only regulate the ability of chromatin to be transcribed but also other DNA-related processes such as recombination, replication, repair, and chromosomal segregation during metaphase^10^. In recent years, a wide range of histone modifications has been described, as well as a limited repertoire of enzymes and proteins that serve to direct the placement (acetylases, methylases, kinases), the removal (deacetylases, demethylases, phosphatases) or the recognition (bromodomains, chromodomains, SANT, PHD fingers, WD40 repeats) of these modifications or “marks”, in what has been called the “histone code”^11^.

One of the most important post transcriptional modifications (PTM) that histone tails undergo is lysine acetylation^12^, a chemical event that is mediated by multiprotein complexes with histone acetyl transferase (HAT) and/or histone deacetylase (HDAC) activities. Many of these complexes also contain bromodomain factors (BDF)^13,14^, proteins with a small domain, the bromodomain (BD), formed by 4 α-helices connected by loops that forms a hydrophobic cavity that recognize acetyl-lysines. Despite components of these signaling pathways have been validated as therapeutic targets for many pathologies, including parasitic diseases^15,16^, the identity of these complexes is just beginning to be revealed and there is little knowledge of their structures.

Recently, Staneva *et al* systematized the analysis of the proteins that compose many putative chromatin regulators, using co-immunoprecipitation (Co-IP) followed by mass spectrometry assays in the process to identify protein networks^17^. They performed Co-IP assays of the 7 BDFs recognizable in the *T. brucei* genome, 9 proteins with acetyl transferase domains and 7 proteins identified as potential HDACs, among others. Another approach to this problem was that of Jones *et al*, who used a proximity labelling methodology to generate an interactomic dataset of BDF5 from *L. mexicana* (*Lmx*BDF5), followed by Co-IP to validate some of the proximity hits as interactions, proposing a novel Conserved Regulator of Kinetoplastid Transcription (CRKT) complex^18^. Also, Vellmer *et al*^19^, while trying to unveil the identity and composition of a novel SNF2 complex involved in H2A.Z deposition, derived in the conclusion that some of the proteins they identified were forming two different HAT complexes, involving *Tb*HAT1 and *Tb*HAT2. In Staneva and Jones’ interactomic datasets, also two distinct MRG (MORF4 related gene) domain-containing proteins were identified. These domains form a structural part of several complexes with HAT and HDAC activity, as described in yeasts and humans^20^. MRG domains seem to act as a “core” or “anchor” to nucleate the protein complex, where multiple proteins compete for a single binding site on the surface of MRG domains. These MRG binding proteins (MRGBPs) interact via intrinsically disordered regions (IDR) that contain an FxLP motif, vital for such interactions^21^. Upon binding, these IDRs acquire an extended conformation, fitting the phenylalanine of the FxLP motif within a small cavity of its partner MRG domain, resting on a conserved arginine^22^. A second segment, more variable, contributes to form a bipartite interaction, also essential to sustain the binding. While the above cited works detected MRG domain containing proteins in *T. brucei* and *L. mexicana*, no MRGBPs were reported there.

On the other hand, recent years have been characterized by an explosion in the development of artificial intelligence algorithms that are contributing to solve previously unsolvable biological problems^23^. One outstanding milestone was the release of the deep learning model AlphaFold2 (AF2), trained to solve the three-dimensional structure of proteins with an unprecedented degree of detail^24^. More recently, after noticing the emerging property of AF2 in predicting protein complexes, AF2-multimer was built and trained using protein complexes as training database, resulting in a higher degree of accuracy^25^.

In this work, we identified the *T. cruzi* homologs of the proteins present in public interactomic datasets, analyzed their sequences, and focused mainly on the search for proteins that interact trough MRG domains. We then generated the structural models of their complexes with AF2-multimer, identifying the presence of two distinct MRG domain-driven assembled complexes. One is the ortholog of the yeast and human TINTIN complexes^26,27^ (that we named *Tc*TINTIN), formed by the protein *Tc*BDF6 and two conserved hypothetical proteins, that we named *Tc*MRGx and *Tc*MRGBP. The second trimeric complex is formed between the MRG domain of *Tc*BDF5 (*Tc*BDF5-MRG), the protein that we named *Tc*BDF5BP (*Tc*BDF5 Binding Protein), and the bromodomain factor *Tc*BDF8, the ortholog of *Lmx*BDF8 described by Jones *et al* in *L. mexicana*^28^. The protein-protein interactions predicted by the quaternary structure models of these two complexes, as well some details about their interactions, were validated using yeast hybrid assays and mutagenesis. Also, we performed molecular dynamic simulations, uncovering a conformational change in *Tc*MRGBP that can explain discrepancies between experimental evidence and AF2-multimer models. Altogether, our results demonstrate that the models produced by AF2-multimer can be used to predict protein-protein interactions in trypanosomatids and are a tool with high predictive value when it comes to inferring protein function, especially when traditional homology-based methods fail.

## Results

Identification of putative trypanosomatid MRG binding proteins (MRGBPs)

To identify the MRG binding proteins, we took as reference the IDs of the interactomic datasets generated by CoIP-MS of *Tb*HAT1 and *Tb*BDF5 produced by Staneva *et al*^29^. *Tb*HAT1 interactome contains the protein Tb927.1.650, which has an HHblits-identifiable MRG domain; along with other 11 significantly enriched proteins, 4 of them annotated as “hypothetical proteins”. By using HHblits^30^, some of them could be identified as constituents of a complex orthologous to the yeast NuA4 (<u>Nu</u>cleosome <u>A</u>cetyltransferase of H<u>4</u>) complex. This is a 13-subunit complex that acts as a chromatin remodeled via its HAT activity and is involved in many processes like transcription and double-strand DNA break repair, among others^31^. However, several of the yeast NuA4 components could not be identified, even among the proteins enriched during reciprocal Co-IPs of other components (*Tb*BDF6 and *Tb*EAF6). Given the high evolutionary distance between trypanosomatids and the model organisms where many chromatin remodeling complexes were described, we considered that some proteins apparently absent from the NuA4 complex could be hidden among the proteins identified as hypothetical in the Co-IPs, being impossible to associate them as orthologs using algorithms such as HHblits due to their high divergence. MRG-binding proteins (MRGBPs) are part of the apparent missing proteins. These proteins are characterized by possessing an intrinsically disordered region (IDR) involved in protein-protein interaction, with a linear motif that possesses an aromatic residue in the first position and a proline in the fourth, which in yeast and humans is an FxLP motif^21^.

On the other hand, *Tb*BDF5 is a protein that has a C-terminal MRG domain. Among the proteins enriched in its interactome, there are two proteins associated with chromatin remodeling: *Tb*HAT2, known to acetylate histone H4 in K2, K5 and K10^32^; and the bromodomain factor *Tb*BDF8. *Tb*BDF5 interactome dataset also contains several hypothetical proteins. One of them is Tb927.9.13320, which possesses an FHA (fork<u>h</u>ead <u>a</u>ssociated) domain, a module involved in phospho-threonine recognition^33^. This protein network was proposed to be forming a complex only found in trypanosomatids, the Conserved Regulators of Kinetoplastid Transcription (CRKT)^18^. No MRGBPs were identified in this interaction network either.

We set out to find the unidentified MRGBPs among the hypothetical proteins by analyzing their sequence, *i*.*e*., by searching for proteins with a predicted IDR that also contain something like a FxLP motif. Once identified, we thought that it would be possible to use the state-of-the-art protein-protein interaction structure prediction algorithm, AF2-multimer^25^, to generate a quaternary structure model of these potential MRGBPs in complex with the identified MRG of the dataset. By looking at the models and their confidence statistics (pLDDT and PAE), we can filter those with bad statistics or that do not match the structural descriptions of MRGBPs. Finally, to validate these interactions we can use a yeast two-hybrid (Y2H) approach^34^ using *T. cruzi* protein sequences. To clone the genes and perform experiments in *T. cruzi*, we used OrthoMCL^35^ to convert by orthology *T. brucei* IDs from the interactomic dataset of *Tb*HAT1 and *Tb*BDF5 to the corresponding ones from *T. cruzi*. **Table 1** and **Table 2** contain the IDs of the orthologs from *T. brucei, T. cruzi* and *L. infantum* reference strains.

**Table 1:** Identified orthologs of the proteins significatively enriched in *Tb*HAT1 CoIP from Staneva *et al*. The identification column contains the protein names or domains identified by Staneva *et al*. \**Tb*YEA2 and *Tb*EAF6 are fused forming a single protein in *L. infantum. Leishmania infantum* orthologous IDs are included because their predicted structures are included in AF2 monomer database.

| *T. brucei* ID | Identification | *T. cruzi* ID | *L. infantum* ID |
| --- | --- | --- | --- |
| Tb927.7.5310 | *TbYEA2* | TcCLB.506825.80 | *LINF_060012900 |
| Tb927.10.14190 | *TbEPL1* | TcCLB.506525.30 | LINF_320005600 |
| Tb927.8.5320 | hypothetical protein, conserved | TcCLB.511217.190 | LINF_160016800 |
| Tb927.1.650 | MRG domain protein | TcCLB.505999.50 | LINF_200005700 |
| Tb927.1.3400 | *TbBDF6* | TcCLB.506529.660 | LINF_120009500 |
| Tb927.11.3430 | hypothetical protein, conserved | TcCLB.507099.100 | LINF_130019500 |
| Tb927.7.4560 | *TbHAT1* | TcCLB.506605.160 | LINF_140006300 |
| Tb927.9.2910 | *TbEAF6* | TcCLB.510565.170 | *LINF_060012900 |
| Tb927.6.1240 | hypothetical protein, conserved | TcCLB.507603.54 | LINF_120005400 |
| Tb927.11.9640 | glycyl-tRNA synthetase, putative | TcCLB.504017.79 | LINF_360047300 |
| Tb927.2.4580 | UNC119 | TcCLB.506979.50 | LINF_270027100 |
| Tb927.10.12040 | mitogen-activated protein kinase 11, putative | TcCLB.510123.20 | LINF_330022100,<br>LINF_190020900 |

**Table 2:**
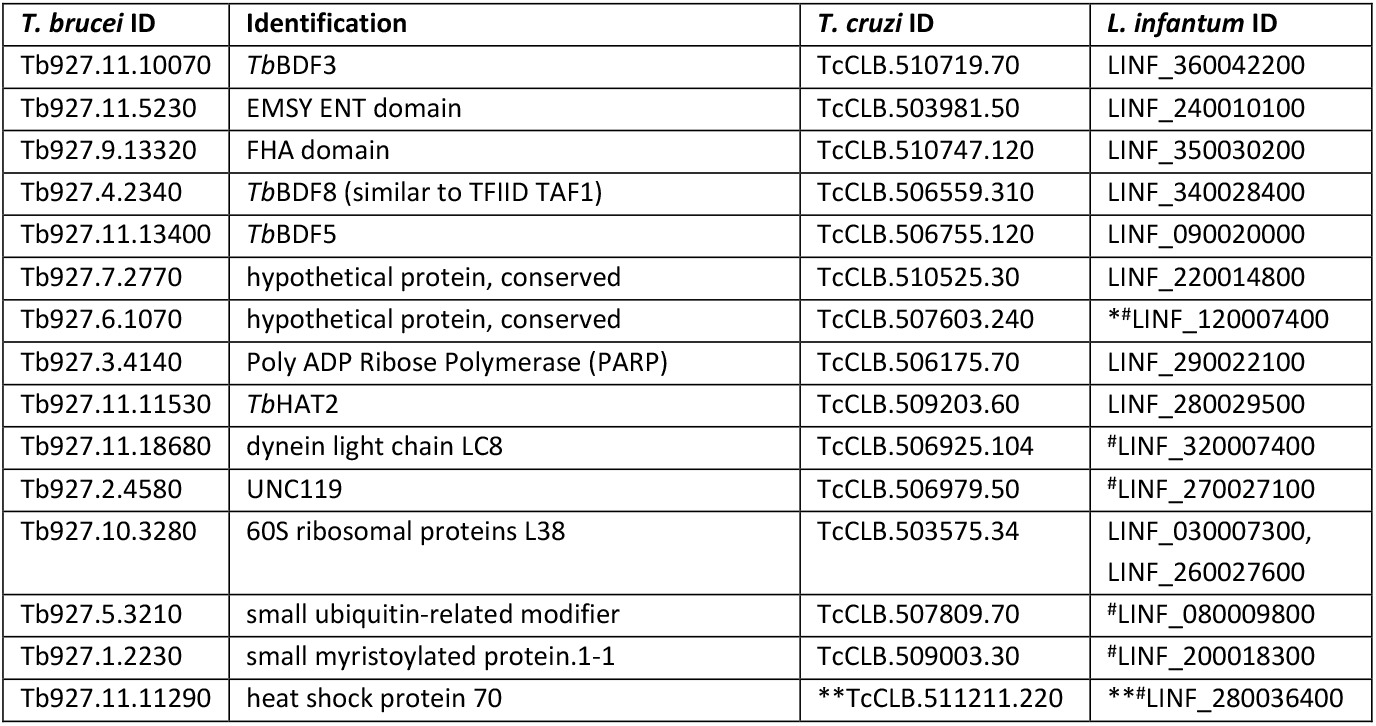
Identified orthologs of the proteins significatively enriched in *Tb*BDF5 CoIP from Staneva *et al* (2021) and in CRKT from Jones *et al* (2022). The identification column contains the protein names or domains identified by Staneva *et al*. *Found by BLAST. **IDs of the syntenic orthologs. *Leishmania infantum* orthologous IDs are included because their predicted structures are included in AF2 monomer database. ^*#*^Not described as part of CRKT.

To identify conserved sequences, we retrieved all trypanosomatid orthologs of each protein using TriTrypDB and aligned each orthology group separately using ClustalO^36^. Because we were interested in IDRs, probability of disorder was predicted with the disorder predictor PrDOS^37^. We also used the pLDDT per residue extracted from the precalculated protein models of *T. cruzi* in AF2 database, since there is a direct correlation between a low pLDDT value and the presence of IDRs^38^. We mapped these annotations on AF2 predicted monomers^39^ and inspected the proteins’ structure models and alignments one by one. In the *Tb*BDF5 CoIP dataset, we identified a conserved Fx[LIV]P motif over an IDR at the N-terminus of *Tc*BDF8 (TcCLB.506559.310) and a conserved [FY]EPP motif at an N-terminal IDR of the FHA domain containing protein TcCLB.510747.120 (**Figure 1a** and **b**, respectively), as the potential interactors of *Tc*BDF5-MRG. BDF8 displays a domain architecture consisting of an N-terminal IDR (residues 1-60) containing the sequence FAIP, a central globular domain of unknown function (DUF) and a C-terminal BD (BD8) (**Figure 1a**). The FHA domain containing protein (TcCLB.510747.120) has three identifiable regions: an IDR N-terminal containing the sequence YEPP, a potential globular DUF with unclear boundaries and an FHA C-terminal (**Figure 1b**). However, we did not find any protein with a conserved motif containing an aromatic aminoacid in the first position and a proline in the fourth among the IDR regions predicted for the proteins on HAT1 interactome. Therefore, we decided to compare their structures with AF2-predicted models of known MRGBPs to search for structural similarities. We found that the hypothetical protein TcCLB.511217.190 has a globular domain followed by an IDR region like that of human MRGBP protein (*Hs*MRGBP). In addition, the region of similarity corresponds to residues involved in the interaction with the human MRG-containing proteins *Hs*MRG15 and *Hs*MRGx (**Figure 1c**), indicating that TcCLB.511217.190 could be a good candidate for binding the protein with MRG domain present in the HAT1 interactome. With this reduced set of candidates, we used AF2-multimer to predict the structures of the pairs TcCLB.505999.50/TcCLB.511217.190, TcCLB.506755.120/TcCLB.510747.120 and TcCLB.506755.120/TcCLB.506559.310. **Table 3** contains the IDs and names we propose for each protein studied in this article, along with proteins already named. The logic by which each protein is named is discussed below. The structural models obtained, the derived structure predictions and the experimental evidence that validates them are described in the following sections.

**Table 3:**
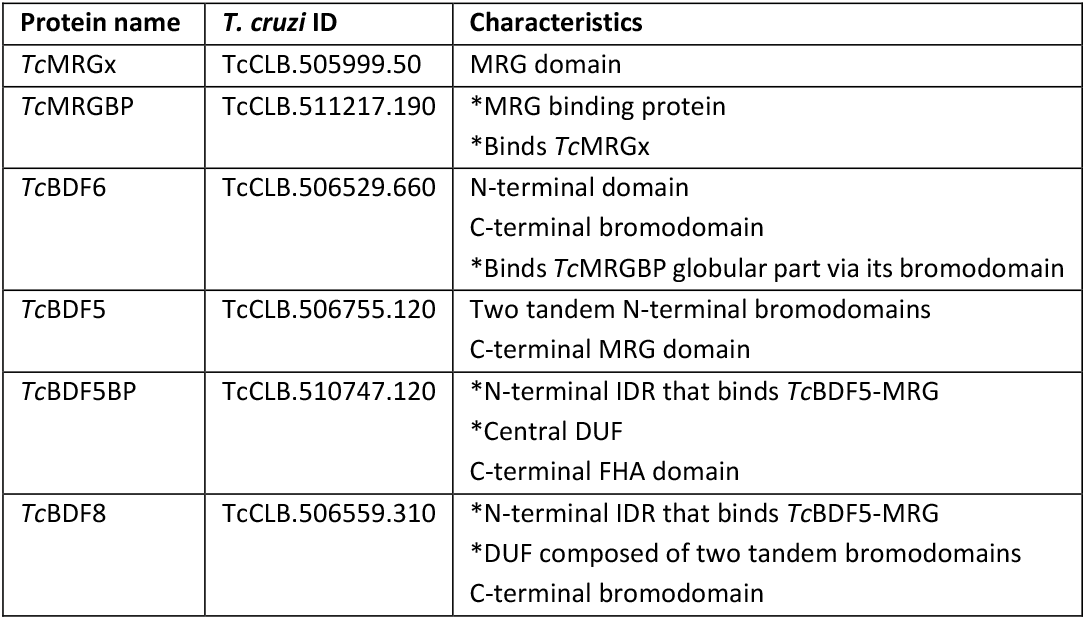
Correspondence between IDs and proposed names of proteins studied in this article. Some characteristics of the proteins are also included. *Described in this study.

| Protein name | *T. cruzi* ID | Characteristics |
| --- | --- | --- |
| *TcMRGx* | TcCLB.505999.50 | MRG domain |
| *TcMRGBP* | TcCLB.511217.190 | *MRG binding protein<br>*Binds *TcMRGx* |
| *TcBDF6* | TcCLB.506529.660 | N-terminal domain<br>C-terminal bromodomain<br>*Binds *TcMRGBP* globular part via its bromodomain |
| *TcBDF5* | TcCLB.506755.120 | Two tandem N-terminal bromodomains<br>C-terminal MRG domain |
| *TcBDF5BP* | TcCLB.510747.120 | *N-terminal IDR that binds *TcBDF5*-MRG<br>*Central DUF<br>C-terminal FHA domain |
| *TcBDF8* | TcCLB.506559.310 | *N-terminal IDR that binds *TcBDF5*-MRG<br>*DUF composed of two tandem bromodomains<br>C-terminal bromodomain |

**Figure 1:**
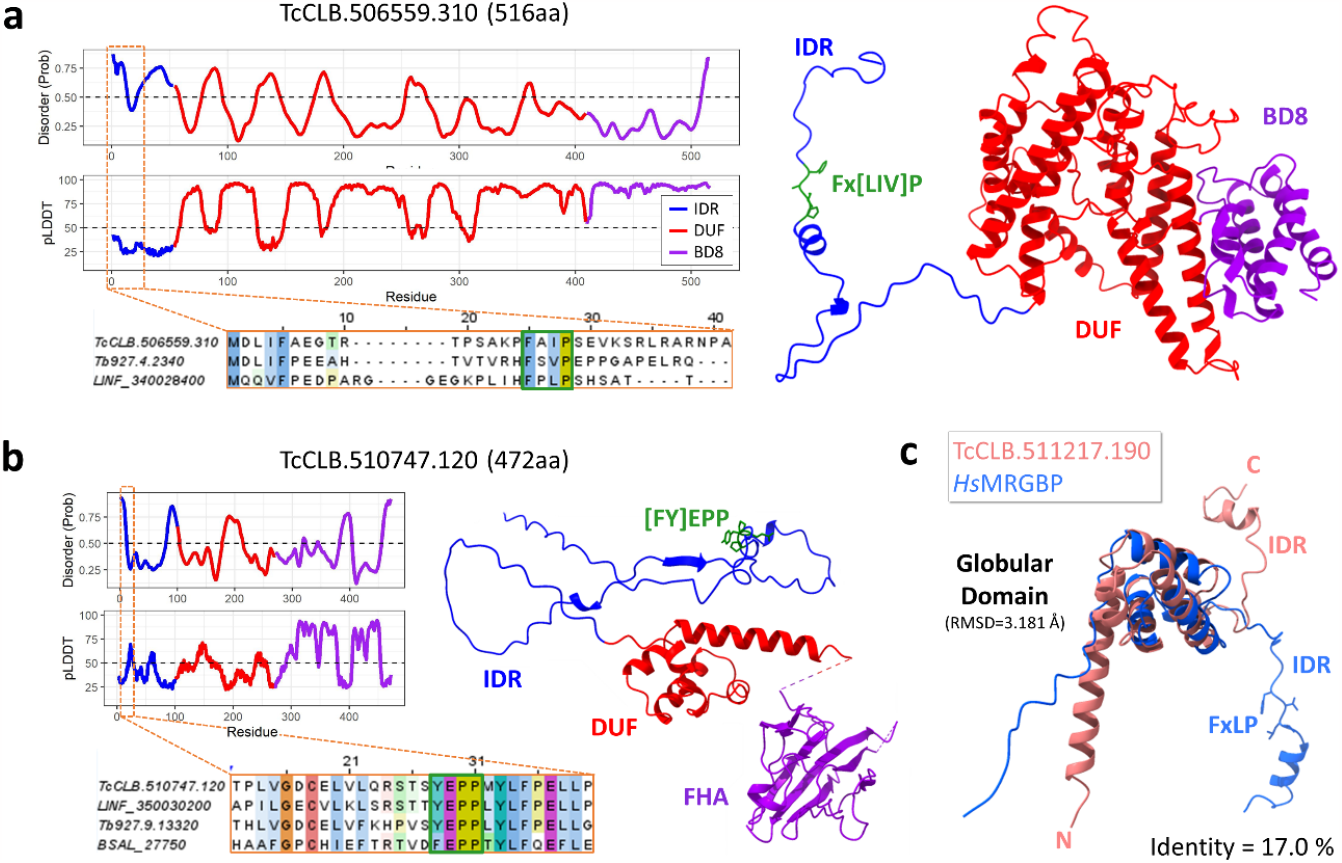
Proteins identified as candidates to be MRGBPs of *T. cruzi*. **(a)** AF2 predicted structure of TcCLB.506559.310 (*Tc*BDF8), disorder probability per residue graph, pLDDT per residue graph and fragment of the multiple sequence alignment for the predicted IDR. Different colors represent the identified domains in both structures and graphs. The dashed orange rectangle points to a zone with low pLDDT and high disorder probability which contains an Fx[LIV]P motif, shown in a green square in the multiple sequence alignment. **(b)** AF2 predicted structure of TcCLB.510746.120 (FHA domain containing protein), disorder probability per residue graph, pLDDT per residue graph and fragment of the multiple sequence alignment for its predicted IDR. As before, each identified domain was colored differently and the dashed orange rectangle indicates the zone in which a conserved [FY]EPP motif was identified. We included the ortholog of *B. saltans*, who contains a phenylalanine in the first position. A disordered loop was removed between the DUF and the FHA domain to enhance visualization. **(c)** Structure comparison between TcCLB.511217.190 and the region of *Hs*MRGBP known to interact with *Hs*MRG15. Both share a globular domain with the same fold, followed by an IDR with a α-helix. The residues corresponding to the FxLP motif of *Hs*MRGBP are shown, along with the sequence identity computed by Needleman-Wunsch algorithm and the RMSD calculation for the globular domain.

### *Tc*TINTIN is a partially conserved eukaryotic complex formed by the sequential assembly of *Tc*MRGx, *Tc*MRGBP and *Tc*BDF6

The structural similarity of *Tc*MRGBP (**Table 3**) with *Hs*MRGBP led us to think that it could be its ortholog. In humans, *Hs*MRGBP is involved in direct binding to the C-terminal MRG domain of *Hs*MRG15 or to the MRG domain of its ortholog *Hs*MRGx, which is smaller, as it lacks the N-terminal chromodomain present in *Hs*MRG15. Furthermore, *Hs*MRGBP can also engage with the bromodomain protein *Hs*BRD8. Together with *Hs*MRG15 (or *Hs*MRGx), they form a complex known as Trimer Independent of NuA4 for Transcription Interactions (TINTIN)^27^, which is also present in *Saccharomyces cerevisiae* formed by *Sc*Eaf3, *Sc*Eaf5 and *Sc*Eaf7^40^. As its name suggests, this complex has 1:1:1 stoichiometry and is part of the NuA4 complex but has transcriptional regulation functions that are independent of NuA4, since it is capable of being assembled independently. Given that *Tb*BDF6 was proposed as an ortholog of *Hs*BRD8^17^, we thought that the set *Tc*MRGx, *Tc*MRGBP and *Tc*BDF6 (**Table 3** and **Figure 2a**) could be forming a TINTIN-like complex in trypanosomatids. The name *Tc*MRGx and not *Tc*MRG15 was chosen because its domain architecture is more like *Hs*MRGx than *Hs*MRG15.

**Figure 2:**
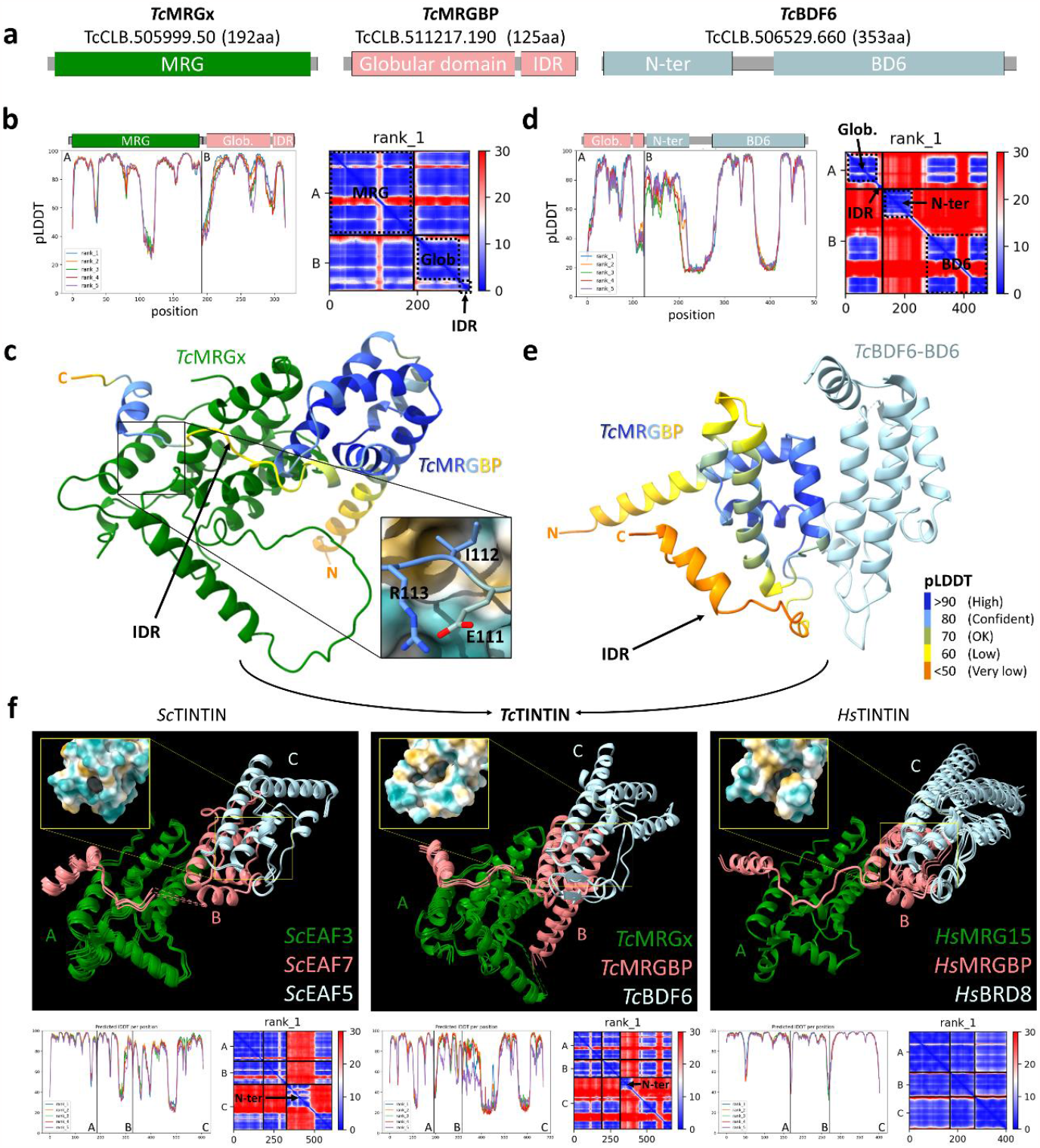
Complex structure prediction of TcTINTIN and its subunits compared to its homologs. **(a)** Domain representations, IDs and length of the proteins involved in the complex. **(b)** Statistics for the AF2-multimer structure prediction of the TcMRGx/TcMRGBP pair. On the left is the pLDDT per residue with the corresponding domains represented over the graph. On the right is the PAE plot with the domain correspondence remarked in dashed squares. **Glob:** Globular domain **(c)** Complex structure prediction of the heterodimer TcMRGx/TcMRGBP. TcMRGx is in green and TcMRGBP is colored by pLDDT (pLDDT color labels are in panel **e**). An arrow points to the predicted IDR in TcMRGBP. The residues of the IDR structure where the FxLP motif is expected to be is enlarged in a rectangle showing the sidechains E111, I112 and R113 of TcMRGBP, with the surface exposed to the solvent of TcMRGx colored by hydrophobicity (yellow: hydrophobic, blue: hydrophilic). **(d)** Statistics for the prediction of the TcMRGBP/TcBDF6 pair. **(e)** Predicted structure for the heterodimer TcMRGBP/TcBDF6. An arrow points to the predicted IDR of TcMRGBP. The N-terminal domain of TcBDF6 was not included in the visualization because it was not involved in the interaction but was included in the prediction. Also, a disordered loop on BD6 was removed to enhance visualization. **(f)** Predicted structures of the heterotrimers of S. cerevisiae TINTIN (ScTINTIN), T. cruzi TINTIN (TcTINTIN) and H. sapiens TINTIN (HsTINTIN) and its statistics (pLDDT and PAE). All five models of each prediction were aligned to the rank_1 model of TcTINTIN in ChimeraX. Disordered loops and the non-interacting N-terminal domains from ScEAF5 and TcBDF6 were removed to facilitate visualization. Notice the lack of low interprotein PAE values in the PAE plot of HsTINTIN. This is because we were computationally unable to include the N-terminal domain of HsBRD8 in the prediction due to a big insertion between it and the bromodomain. The curved black arrows between panels **c** and **e** indicate that the combination TcMRG/TcMRGBP/TcBDF6 forms TcTINTIN. The hydrophobic pockets of ScEAF5, TcBDF6 and HsBRD8 are shown in an inset for each trimer.

**Figure 2b** shows the statistics for the AF2-multimer prediction of *Tc*MRGx/*Tc*MRGBP pair. The predicted aligned error (PAE) plot indicates a low predicted error between the relative positions of the amino acids of both proteins (upper right and lower left quadrants), so it is possible to consider the generated model (**Figure 2c**) as the structural prediction of this dimer. The pLDDT plot, which represents the confidence per-residue, indicates good confidence in the individual domains generated, with values below 60 only for some loops connecting α-helices. Values greater than 80 can even be observed at IDR zone that contacts *Tc*MRGx, indicating that AF2 is confident in its local structure. The mode of interaction is similar to the bipartite interaction observed for the NMR structure of the *Hs*MRG15/*Hs*MRGBP pair (PDB: 2N1D)^21^, involving multiple hydrophobic amino acids, both in the globular region, and in the IDR (**Figure S1**). Regarding the residues where the FxLP motif should be, the I112, partially conserved among trypanosomatid orthologs, is found in *Tc*MRGBP, while *Tc*MRGx does not display any clear hydrophobic pocket over its conserved arginine R138 (on which I112 is resting). This suggests that this interaction may not be as important as in other MRGBPs (see below).

The statistics and the model for the prediction of the *Tc*MRGBP/*Tc*BDF6 pair are shown in **Figure 2d** and **Figure 2e**, respectively. The PAE plot shows values with low relative error only between the residue pairs involving the globular domain of *Tc*MRGBP and the BD of *Tc*BDF6, suggesting that the IDR of *Tc*MRGBP and the N-terminal domain of *Tc*BDF6 are not involved in the interaction. Consistently, a decrease in pLDDT values over the IDR of *Tc*MRGBP can also be seen on this model when compared to the model of *Tc*MRGx/*Tc*MRGBP dimer from **Figure 2c**, suggesting that the IDR is only involved in the interaction with *Tc*MRGx, and not with *Tc*BDF6. A low pLDDT value was also observed over the IDR residues of *Tc*MRGBP monomer, indicating that *Tc*MRGBP may be acquiring conformational stability only after binding to *Tc*MRGx. There is also a slight decrease in pLDDT values of other *Tc*MRGBP residues that contact *Tc*MRGx but not *Tc*BDF6.

Finally, we predicted the structure of the three-component complex (*Tc*MRGx/*Tc*MRGBP/*Tc*BDF6), which we will refer to as *Tc*TINTIN from now on. We did the same for the proteins *Sc*EAF3, *Sc*EAF7 and *Sc*EAF5, which together form the yeast TINTIN complex (*Sc*TINTIN); and with the proteins *Hs*MRG15, *Hs*MRGBP and *Hs*BRD8, which assemble *Hs*TINTIN. A comparison between these predictions can be seen in **Figure 2f**. The PAE of the three complexes show good confidence between the relative positions of the atoms of all the domains of the three proteins, except for the N-terminal domains of *Sc*EAF5 in *Sc*TINTIN and *Tc*BDF6 in *Tc*TINTIN. This suggests that the N-terminal domains are not involved in the interaction. Due to video memory size limitations, we restricted the length of *Hs*BRD8 only to its BD residues, since it has a long insertion between the BD and its N-terminal domain, the latter being homologous to the N-terminal domains of *Tc*BDF6 and ScEAF5 (**Figure S2**). So, it was not included in the prediction of *Hs*TINTIN, but we would expect to see a similar chart with low PAE values relative to the rest of the domains if it could be included. A good general correlation between high values of pLDDT and low values of intraprotein PAE is also observed, indicating good packing of the individual domains. From the structural point of view, the models present a similar spatial arrangement of the proteins that compose them. ScEAF7, TcMRGBP and HsMRGBP show the classic bipartite interaction described for MRGBPs, extending their IDR over the surface of their partner MRG domain, also binding via their globular domains. Meanwhile, *Sc*EAF5, *Tc*BDF6 and *Hs*BRD8 mediate their interaction with *Sc*EAF7, *Tc*MRGBP and *Hs*MRGBP, respectively, via two of their α-helices, exposing a hydrophobic pocket to the solvent that can potentially recognize acetylated lysines.

Although these models are consistent and clearly describe homologous structures, AF2-multimer has not yet been widely validated, nor is there experimental evidence indicating that its predicted quaternary structures can be assumed legitimate in trypanosomatids. Therefore, we set out to validate the formation of these complexes *in vivo* using the Y2H methodology previously used in our lab to test protein-protein interactions in multicomponent complexes^34^. In this methodology, protein-protein interaction is measured in yeasts by the expression level of reporter genes which confer the ability to grow in absence of essential molecules, together with the expression of β*-gal* whose expression can be easily seen through β-galactosidase-mediated colorimetric reactions^41^.

Initially, we tested *Tc*MRGx/*Tc*MRGBP and *Tc*MRGBP/*Tc*BDF6 interactions. As expected, yeast clones expressing the *Tc*MRGx/*Tc*MRGBP interaction pair showed β-galactosidase activity (**Figure 3a**). The blue color produced after X-Gal hydrolysis could be observed even one-hour post-incubation, suggesting that *Tc*MRGx/*Tc*MRGBP are strong interactors. No *Tc*MRGx, neither *Tc*MRGBP alone were able to induce reporter genes’ expression (autoactivating controls). However, no β-galactosidase activity was seen for *Tc*MRGBP/*Tc*BDF6 pair, an unexpected result given the high confidence observed in the statistics of the models produced by AF2-multimer for this pair of proteins (**Figure 2**). We thought that it was because *Tc*MRGBP/*Tc*BDF6 interaction was too weak to see with β-gal as reporter. Therefore, we tested *HIS3* reporter, which has a stronger promoter, but the spot growth assays performed did not show interaction phenotype either (**Figure S3**). We also checked the presence of the proteins by western blot to discard any defect in protein expression in yeasts (**Figure S4**). Finally, we reanalyzed the interaction models, and realized that the motility of the free IDR of *Tc*MRGBP could only be retained after binding to *Tc*MRGx. Due to this, in a scenario where *Tc*MRGx is absent, the IDR chaotic movement could prevent the formation of a stable complex with *Tc*BDF6. But if *Tc*MRGx is present in the system, the IDR would stabilize (maybe inducing a general conformational change), allowing the assembly of the complete complex. If this is true, it would mean that *Tc*TINTIN should assemble in a sequential fashion.

**Figure 3:**
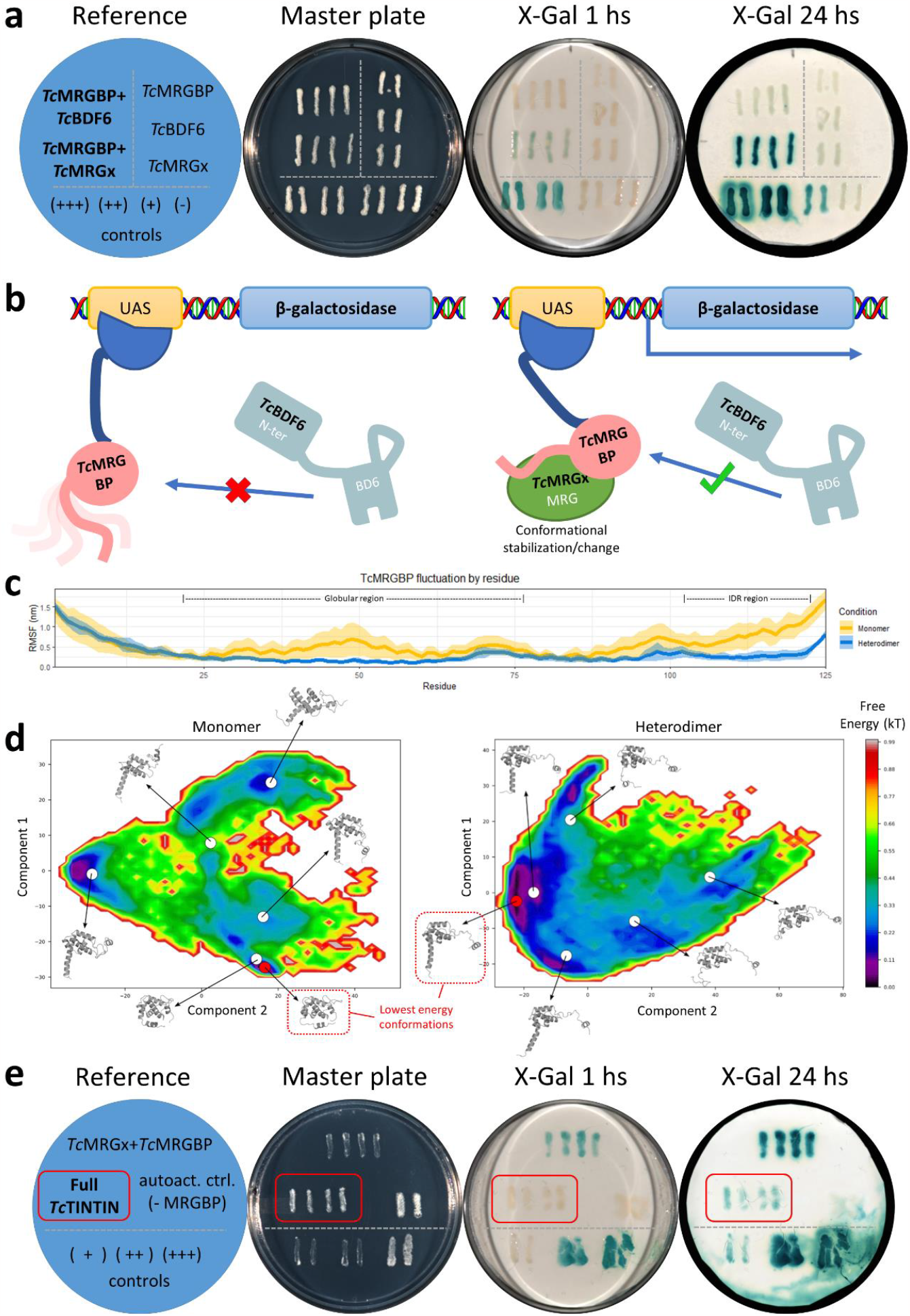
Y2H and Y3H assays validate the formation of *Tc*TINTIN *in vivo* and suggests that is assembled sequentially: a possible mechanism derived from molecular dynamics. **(a)** Y2H assay to test the interaction pairs *Tc*MRG/*Tc*MRGBP and *Tc*MRGBP/*Tc*BDF6 using β-galactosidase as reporter gene. The blue circle has a reference to the yeast strains streaked on the master plate (SC-Leu-Trp). Control strains were streaked below the horizontal line, autoactivation controls were streaked at the right of the vertical line and the actual interaction tests to the left. **(b)** Cartoon of *Tc*TINTIN sequential assembly hypothesis. Left: *Tc*BDF6 cannot interact with *Tc*MRGBP in absence of *Tc*MRGx. Right: *Tc*MRGBP binds to *Tc*MRGx and changes conformation, activating the interaction with *Tc*BDF6 and transcribing the reporter gene. **(c)** Fluctuations on each residue of *Tc*MRGBP using RMSF by residue plot derived from molecular dynamics (MD) simulations. RMSF values of *Tc*MRGBP simulated as a monomer is in yellow and in blue as a heterodimer. Thick lines are the mean RMSF, and transparent colors are the standard deviations. **(d)** WHAM plots derived from MD simulations. Free-energy maps were calculated using RMSD as reaction coordinate and medoid structures were obtained with agglomerative ward algorithm. Lowest energy conformations are highlighted in red. Left plot corresponds to *Tc*MRGBP simulated as a monomer and right plot as heterodimer. **(e)** Results from the Y3H experiment using β-galactosidase as reporter gene. The reference indicates the yeast strains that were streaked on the master plate (SC-Leu-Trp-Ura). The negative interaction control strain (-) was not included because it does not growth in absence of Ura, used to select GFP-*Tc*MRGx expression plasmid. A red rectangle indicates the yeast clones that express the three proteins from panel b-right, showing β-galactosidase signal at 24 hours.

To further explore this hypothesis, we conducted molecular dynamics (MD) simulations of TcMRGBP under two conditions: one with TcMRGBP isolated as a monomer, and the other with TcMRGBP engaged with TcMRGx as a heterodimer. Each condition was simulated in triplicate, and several metrics were calculated and analyzed alongside the structural coordinates. **Figure 3c** displays the fluctuation of TcMRGBP residues during the simulations, presented as RMSF values by residue. The results indicate significantly higher fluctuations of the IDR residues in the monomeric condition compared to the heterodimeric one.

Additionally, significantly higher fluctuations were observed in the globular region. These findings further support the hypothesis that *Tc*MRGx acts as a structural stabilizer of *Tc*MRGBP. However, RMSF values only provide information about the average fluctuations during simulations and do not explain why *Tc*BDF6 is unable to bind to *Tc*MRGBP in its monomeric form. To address this issue, we analyzed the combined structures of each condition and calculated the free-energy map using the WHAM method ^42^ (**Figure 3d**). Using these maps, we identified the free energy minima of both systems, which revealed two distinct conformational states of *Tc*MRGBP. The first state is compact, with low solvent accessibility and is only present at the lowest free energy state when *Tc*MRGBP is simulated as a monomer. The second state is an extended conformational state present in both systems, but it corresponds to the lowest free energy state only when *Tc*MRGBP is simulated in a complex with *Tc*MRGx. Together, these findings support the idea of structural stabilization induced on *Tc*MRGBP upon binding to *Tc*MRGx, suggesting that it locks *Tc*MRGBP in a stable, extended conformational state, which ultimately allows for a stable interaction with *Tc*BDF6. A comprehensive analysis of the MD simulations can be found in the Supporting Information.

To test this hypothesis (**Figure 3b**) in the laboratory, we generated a third plasmid coding for *Tc*MRGx fused to GFP and introduced it into the strain expressing *Tc*MRGBP/*Tc*BDF6 interaction pair to perform a yeast-three hybrid (Y3H) experiment. Correct expression of GFP-*Tc*MRGx in the nucleus was verified by fluorescence microscopy (**Figure S5**). Fluorescent colonies were streaked into a master plate to evaluate their β-galactosidase activity, together with colonies from the strain expressing the pair *Tc*MRGx/*Tc*MRGBP to compare (**Figure 3c**). After 24 hours, clones expressing full *Tc*TINTIN (*Tc*MRGx/GFP-*Tc*MRGBP/*Tc*BDF6) showed β-galactosidase activity, while the autoactivation controls did not show it at any time. To confirm that the conformation of the *Tc*MRGBP/GFP-*Tc*MRGx/*Tc*BDF6 trimer is not due to a spurious interaction of the GFP protein with *Tc*MRGBP and *Tc*BDF6, a yeast expressing *Tc*MRGBP/GFP/*Tc*BDF6 was included among the controls (**Figures S6 and S7**).

These results do not only validate the formation of *Tc*TINTIN complex but also gives information about the dynamic of its assembly. The sequential assembly observed in this study could potentially enable finer control over the formation of *Tc*TINTIN. By regulating the formation of the complex at specific points in the assembly process, the cell could ensure that *Tc*TINTIN only forms under certain conditions or in response to certain signals. This could help prevent premature complex formation or formation in inappropriate contexts.

### The MRG domain of *Tc*BDF5 forms an heterotrimer only observed in trypanosomatids that displays novel MRGBPs interaction modes

Structural comparison between the MRG domain of *Tc*BDF5 and *Hs*MRG15 shows one mayor insertion of approximately 50 aminoacids between α helices 1 and 2 (Loop1-2), and two minor insertions, one of which is also present in *Tc*MRGx (Loop4-5), but not the other (Loop5-6) (**Figure S8**). Such differences suggest that *Tc*BDF5-MRG is a paralog of *Hs*MRG15 and *Tc*MRGx. We used the process described above for the predicted *Tc*BDF5-MRG interactors. A representation of the domain architecture of *Tc*BDF5 and the proteins predicted as potential interactors of its C-terminal MRG domain is shown in **Figure 4a**. We made several structural predictions for the pair constituted by *Tc*BDF5 and the FHA domain-containing protein that we named *Tc*BDF5BP (**Table 3**), identifying low relative PAE values only between the amino acids of the MRG domain of *Tc*BDF5 and the IDR region of *Tc*BDF5BP (**Figure 4b**). Comparing the pLDDT values for the IDR region observed in *Tc*BDF5BP monomer of **Figure 1b** and those of the heterodimer, we saw an increase in the maximum values from ∼60 in the former to ∼95 in the latter, pointing to a greater confidence in the prediction of its local structure in the heterodimer and a structural stabilization. The binding of this IDR shows the typical extended conformation, positioning the aromatic residue of the [FY]EPP motif inside a cavity formed on the surface of its partner MRG domain *Tc*BDF5-MRG (**Figure 4c**), that is why we decided to name this protein as *Tc*BDF5BP (*Tc*BDF5 Binding Protein). Nonetheless, it also displays a novel way of interaction for MRGBPs, with the IDR in an extended-turn-extended conformation. The models indicate that both DUF and FHA domain are not involved in the formation of the heterodimer (**Figure S9**).

**Figure 4:**
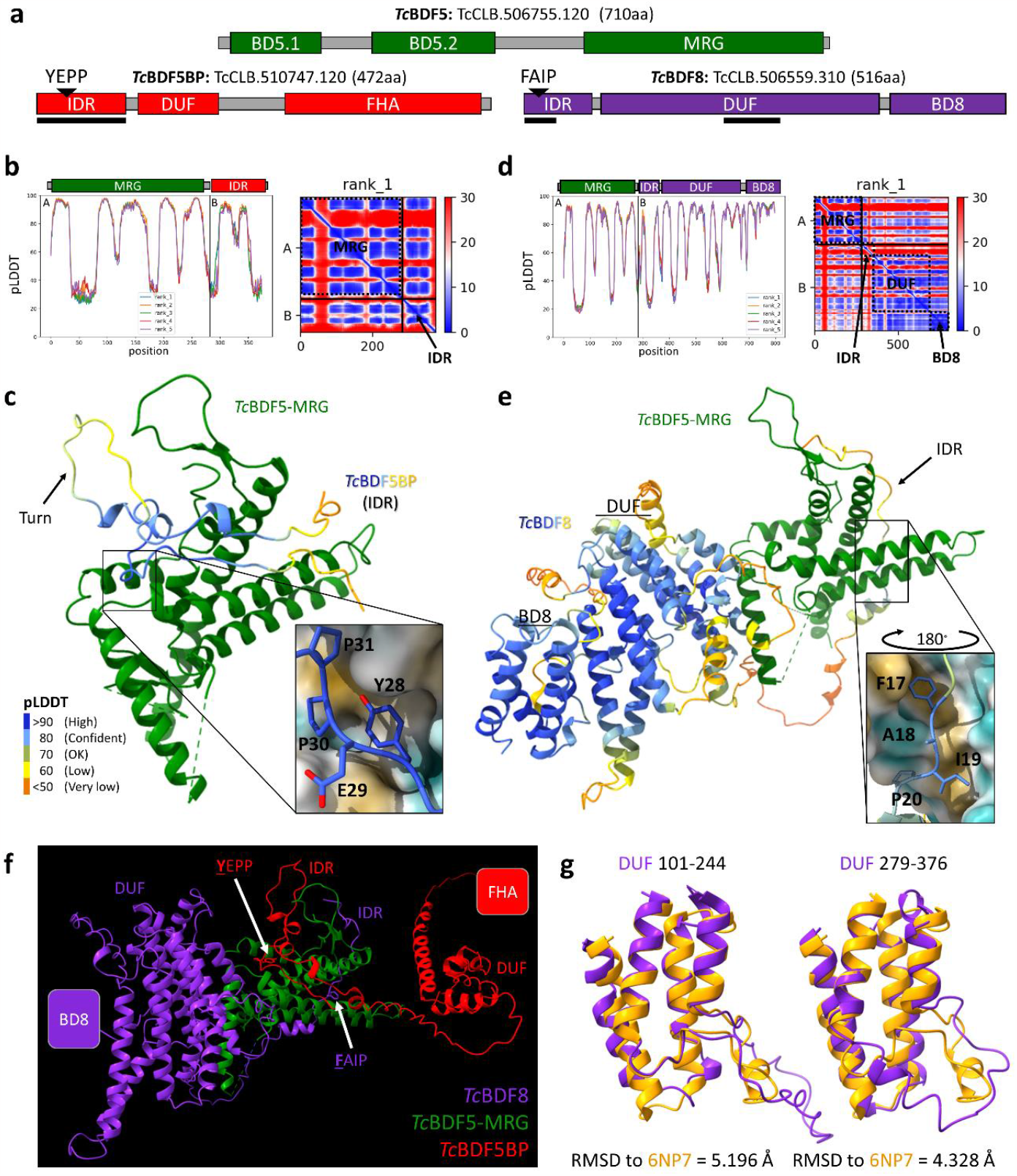
Predicted complex formed via the MRG domain of *Tc*BDF5. **(a)** Domain representation, IDs and length of the proteins involved in the complex. Positions of the YEPP sequence corresponding to the [FY]EPP motif and FAIP sequence corresponding to the Fx[LIV]P motif are indicated with a triangle over the IDRs of *Tc*BDF5BP and *Tc*BDF8, respectively. Solid black lines indicate the interaction regions with *Tc*BDF5-MRG predicted by AF2-multimer. FHA: Forkhead associated domain. DUF: Domain of unknown function. BD8: Bromodomain of *Tc*BDF8. **(b)** Model confidence statistics for the *Tc*BDF5-MRG/*Tc*BDF5BP-IDR heterodimer. **(c)** Model of the complex formed between the IDR of *Tc*BDF5BP and the MRG of *Tc*BDF5. *Tc*BDF5 is colored green and *Tc*BDF5BP is colored by pLDDT. The mode of interaction of the YEPP sequence is seen in the box, which is analogous to the FxLP motif observed in other MRG-domain partners. **(d)** Model confidence statistics for the *Tc*BDF5-MRG/*Tc*BDF8 heterodimer. **(e)** Structure prediction of the *Tc*BDF5-MRG/*Tc*BDF8 heterodimer. *Tc*BDF5 is colored green and *Tc*BDF8 is colored by pLDDT (pLDDT color labels are in panel **c**). An arrow points to the IDR of *Tc*BDF8. The mode of interaction of the FAIP sequence is seen in the box with a 180° rotated point of view, with the surface exposed to the solvent of *Tc*BDF5-MRG colored by hydrophobicity. A black arrow points to the IDR domain of *Tc*BDF8. **(f)** Model of the trimer formed by *Tc*BDF8/*Tc*BDF5/*Tc*BDF5BP, predicted with AF2-multimer. **(g)** Structure alignment between the two bromodomain-like structures of *Tc*BDF8-DUF in purple and the crystal structure of the BD of *Tc*BDF2 in light brown (PDB: 6NP7). RMSD calculations are shown below.

In an attempt to find potential homologs of *Tc*BDF5BP, we constructed a phylogenetic tree using the seed sequences of Pfam^43^ FHA protein family (PF00498) and the FHA domain sequences of BDF5BP from *T. cruzi, T. brucei* and *L. infantum* (**Figure S10**). In this tree, the FHA clusters in a clade of proteins with similar size and domain architecture to that of *Tc*BDF5BP, *i*.*e*., around 500 aminoacids, an N-terminal IDR, a central globular domain, and a C-terminal FHA domain. They even had a coiled-coil between their central globular domain and the FHA, also observed between the DUF and FHA domains of *Tc*BDF5BP. Among them are *Hs*MCRS1 and *Dm*Rcd5 (*Drosophila melanogaster* Rcd5), which are part of various chromatin remodeling complexes. In particular, they are part of the NSL (Non-Specific Lethal) complex, with a fundamental role in dosage compensation of X chromosome through the acetylation of histone H4 in lysines 5, 8 and 16^44^. This complex shares its HAT enzyme (MOF) with another complex named MSL (Male-Specific Lethal). MOF modifies its substrate specificity when it is in MSL, losing the ability to acetylate lysines 5 and 8. Contrary to NSL, MSL contains an MRG protein, paralogous to MRG15, known as MSL-3^45^. As far as we know, there is no report about *Hs*MCRS1 or *Dm*Rcd5 binding any MRG domain.

For the predictions of the *Tc*BDF5/*Tc*BDF8 pair, we find models describing interaction not only through the predicted IDR of *Tc*BDF8, but also involving part of the DUF (**Figure 4d** and **e**), involved in what seems to be a bipartite interaction, one of the characteristics described for known MRGBPs^21^. As occurred with *Tc*BDF5BP, the pLDDT values for the IDR in *Tc*BDF8 monomer (**Figure 1a**) are lower (∼40) than those reached in the dimer (∼85), indicating an improvement in the confidence of the packaging of these residues when the MRG of BDF5 is present in the prediction. In particular, the IDR peaks in pLDDT over the region containing the Fx[LIV]P motif. Structurally, the phenylalanine deepens into a hydrophobic cavity formed between two α-helices of *Tc*BDF5-MRG. This hydrophobic pocket has never been reported as a binding site for the conserved motif of any MRGBP. The other region of interaction occurs on the opposite side of the MRG domain, where several DUF α-helices wrap around one *Tc*BDF5-MRG α-helix. Taken together, this bipartite interaction of *Tc*BDF8 does not occupy the same cavities involved in *Tc*BDF5BP binding. Typically, several MRGBPs compete for the same binding site on the MRG^46^, assembling different complexes depending on the context. In this case, AF2-multimer prediction using the three proteins suggest that *Tc*BDF5 is forming a trimer through its MRG domain with *Tc*BDF5BP and *Tc*BDF8 (**Figure 4f**).

While reviewing the structures, we noticed a structural similarity between the folds of the α-helices present in *Tc*BDF8-DUF domain with the bromodomain fold (**Figure 4g**). We made several structural alignments with different BD crystal structures and concluded that DUF is composed of two tandem bromodomain-like structures packaged into one globular domain. The only residues clearly conserved in *Tc*BDF8-DUF are mainly hydrophobic and located in opposed pairs at the core of the bromodomain-like structures, as if they had only a structural function in keeping the domain folded (**Figure S11a**). We saw no signs of conserved residues or hydrophobic pocket potentially involved acetyl-lysine recognition. On the contrary, this do not happen with the C-terminal bromodomain (BD8), which, although it does not have an essential asparagine involved in acetyl-lysine recognition, it has multiple clearly conserved hydrophobic residues on a small pocket (**Figure S11b**). Therefore, even though the bromodomain-like structures of DUF would have mainly a structural function, we cannot rule out the possibility that BD8 recognizes acetylated lysines.

To perform the Y2H assays, we fragmented *Tc*BDF5BP and *Tc*BDF8 to compare eventual positive interactions with the predicted heterodimers. We generated truncated versions of *Tc*BDF5BP: one that only contains the IDR (IDR), another that contains the rest but not the IDR (NoIDR) and a third with the complete CDS (Full). To avoid the potential autoinduction caused by the BDs of *Tc*BDF5 binding to chromatin, we performed the tests only using its MRG domain. The results for the pair *Tc*BDF5-MRG/*Tc*BDF5BP are shown in **Figure 5a**. As predicted by the models, we only found β-galactosidase activity in yeasts that contain the MRG domain of *Tc*BDF5 together with the *Tc*BDF5BP versions that contain the IDR (Full and IDR on the figure). The blue color could be detected after one hour of incubation, suggesting a strong interaction. For *Tc*BDF8 we generated truncated versions: only its IDR (IDR), only DUF (DUF), only BD8 (BD8) and the complete CDS (Full). Again, we tested them against the MRG domain of *Tc*BDF5 (**Figure 5b**). We were expecting to see at least weak interactions for *Tc*BDF5-MRG tested with IDR or DUF, along with no interaction with BD8 and a strong one with the full protein. Instead, we saw β-galactosidase activity only when the full protein was expressed after 24 hours of incubation. This suggests that the heterodimer can only be formed if both parts of the predicted bipartite interaction (IDR and DUF) are present in the CDS of *Tc*BDF8. Mutagenesis analysis of *Hs*Pf1, a human MRG binding protein that maintains a bipartite interaction with HsMRG15, has shown similar results; mutations that affect only one side of the bipartite interaction completely disrupted the interaction^21^.

**Figure 5:**
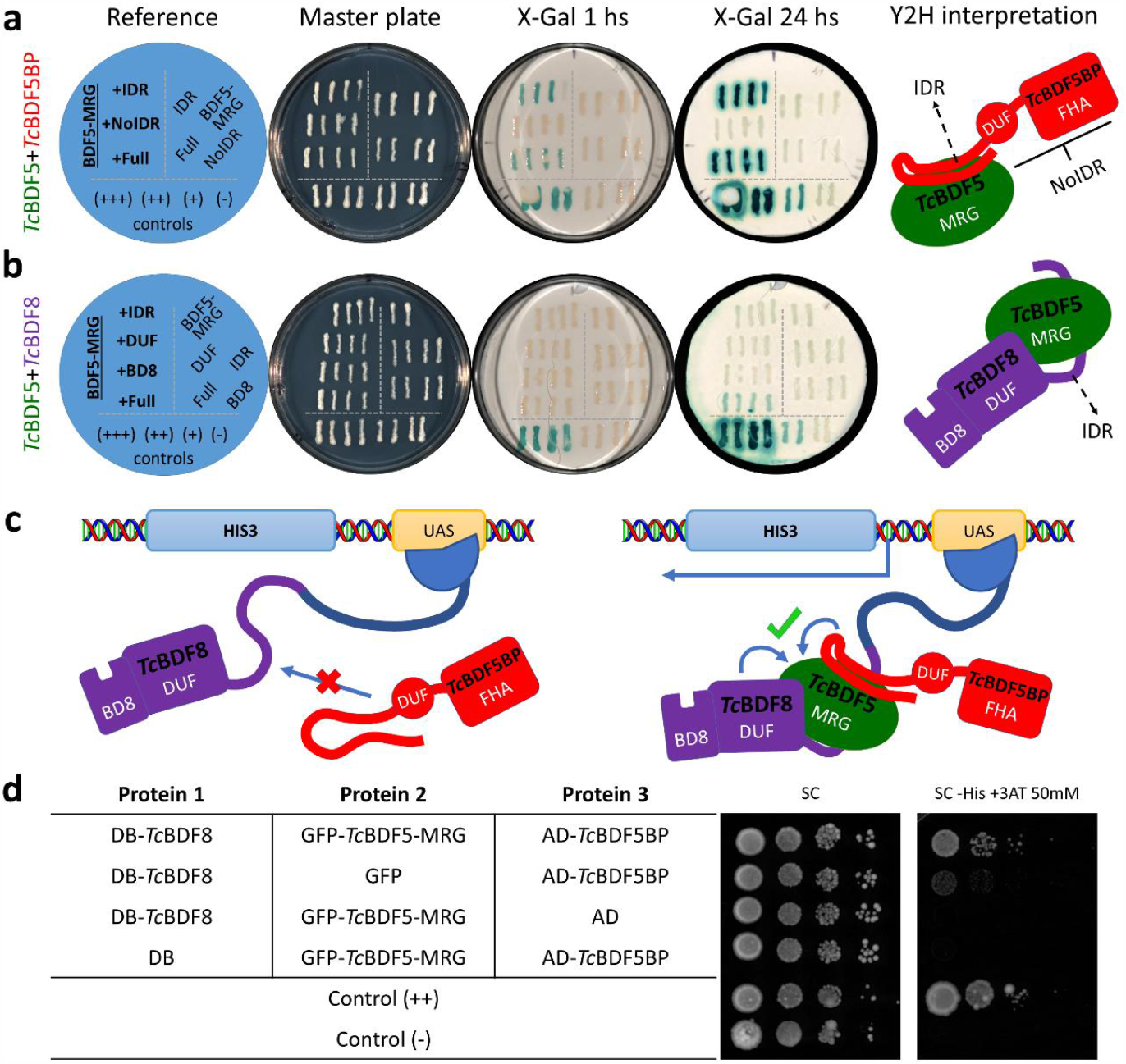
Y2H and Y3H assays validate the formation of the predicted *Tc*BDF5-MRG mediated trimer. **(a)** Y2H assay between different versions of *Tc*BDF5BP and the MRG domain of *Tc*BDF5 β-galactosidase as reporter gene. Control strains are shown below the horizontal line, autoactivation strains were streaked to the right of the vertical line and interaction tests strains expressing *Tc*BDF5-MRG and different versions of *Tc*BDF5BP are to the left of the vertical line. An interpretation of the assay is show as a cartoon to the right. IDR: CDS containing just the IDR of *Tc*BDF5BP. NoIDR: CDS containing *Tc*BDF5BP without its IDR. Full: Full length of *Tc*BDF5BP. **(b)** Y2H between different versions of *Tc*BDF8 and the MRG domain of *Tc*BDF5. As before, a similar distribution of the strains was applied. IDR: CDS containing just the IDR of *Tc*BDF8. DUF: CDS containing just the DUF of *Tc*BDF8. BD8: CDS containing just the BD of *Tc*BDF8. Full: Full length of *Tc*BDF8. **(c)** Depiction of the Y3H assay set to validate the simultaneous binding of *Tc*BDF8 and *Tc*BDF5BP to different surfaces of *Tc*BDF5-MRG. **(d)** Results of the Y3H assay performed as spot growth assay using *HIS3* as reporter gene. The proteins co-expressed in each strain are shown in the table. Serial dilutions of each strain were deposited from left to right as drops in SC (growth control) and SC -His +3AT (test) and incubated for 6 days at 30 °C. DB: GAL4 DNA binding domain. AD: GAL4 activation domain.

**Figure 6:**
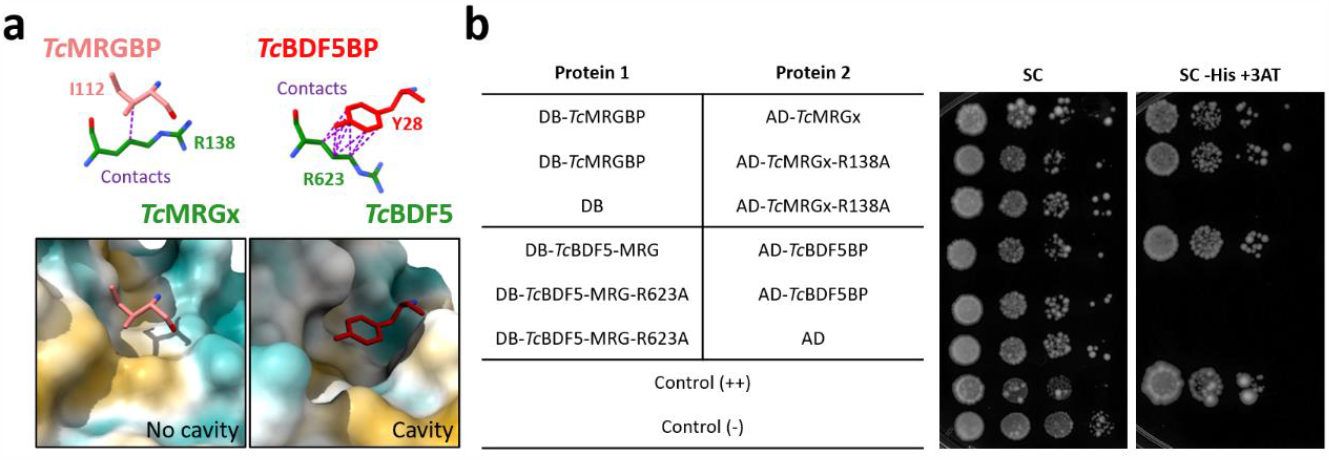
Number of MRGBPs atoms in contact with the conserved arginine of its partner MRG correlates with the loss of interaction when mutated by alanine. **(a)** The I112 of *Tc*MRGBP that is in the position where the aromatic residue of the conserved FxLP motif should be, shows only has a single contact with the conserved arginine R138 of *Tc*MRGx in the AF2-multimer prediction. In contrast, the aromatic residue Y28 of the [FY]EPP motif of *Tc*BDF5BP is in close contact (9 atom to atom contacts) with the conserved arginine R623 of *Tc*BDF5. Note: R623 is the conserved arginine in position 623 of Dm28c *Tc*BDF5, that is R634 for CL Brener (see materials and methods). **(b)** Spot growth assay of MaV203 strains expressing a combination of DB, AD, DB-*Tc*MRGBP, DB-*Tc*BDF5-MRG, DB-*Tc*BDF5-MRG-R623A, AD-*Tc*MRGx, AD-*Tc*MRGx-R138A and AD-*Tc*BDF5BP. The table on the left indicates the combinations tested in each row. Positive and negative interaction controls were also included. Dilutions of OD_600_=1.000, 0.100, 0.010 and 0.001 were spotted on solid SC medium or solid SC medium without histidine (-His) supplemented with 50mM 3-AT to titrate basal HIS3 expression. Cells were incubated at 30 °C for 8 days.

However, these observations alone are insufficient to prove that the binding surfaces on *Tc*BDF5-MRG of *Tc*BDF8 and *Tc*BDF5BP are different. It is possible that they bind to the same interface, as they share a motif with an aromatic amino acid in the first position and a proline in the fourth. Additionally, AF2 predictions of *Tc*BDF8/*Tc*BDF5 dimer may be incorrect. To address these concerns, we designed a new Y3H assay in which the expression of *HIS3* as a reporter gene can only be induced if the interaction surfaces are different (**Figure 5c**).

In this experimental setup, *Tc*BDF8 and *Tc*BDF5BP were fused to the AD and DB domains of the Y2H system. If the predictions of AF2-multimer are correct, the proteins alone would not be able to interact with each other and would only activate *HIS3* transcription if *Tc*BDF5-MRG is present in the system, acting as a bridge between these two protein fusions. For the Y3H assay, we used *Tc*BDF5-MRG fused to GFP, or only GFP in a third vector. The results of this experiment are shown in **Figure 5d**. In the spot growth assay, we observed that the condition containing the complete Y3H system showed the highest cell growth. It grew even at the highest dilutions and developed larger colonies than any of the controls.

In summary, these observations confirm the formation of a complex composed of *Tc*BDF8/*Tc*BDF5/*Tc*BDF5BP, which likely exists in a 1:1:1 stoichiometry. *Tc*BDF8 and *Tc*BDF5BP directly interact with *Tc*BDF5 through its MRG domain on different surfaces. Meanwhile, AF2-multimer models do not predict homodimerization of any of these proteins.

### The number of predicted contacts with the MRG conserved arginine correlates with its importance in maintaining MRGBPs interaction

As discussed above, *Tc*MRGBP do not have any sequence in its IDR region that fit the FxLP motif described in other organisms (**Figure 2c**). Instead, it has a partially conserved isoleucine that contacts loosely with the conserved arginine R138 of *Tc*MRGx (**Figure *6*a**). Furthermore, contrary to what is described for other MRGs^47^, the surface exposed to the solvent around R138 has no clear hydrophobic pocket. Meanwhile, *Tc*BDF5BP has a conserved tyrosine (Y28) in its [FY]EPP motif that closely contacts the aliphatic portion of the conserved Arg R623 of *Tc*BDF5. Also, a clear cavity is present in this region, in which Y28 deepens. By comparison, *Tc*MRGx-R138 has 1 predicted atom to atom contact with *Tc*MRGBP-I112, while *Tc*BDF5-R623 shares 9 with *Tc*BDF5BP-Y28 (**Figure *6*a**).

In order to see if these structural observations were correlated to the strength of the interaction established between the MRGs and their corresponding MRGBPs, we mutated the conserved Arg in both MRG domains to Ala and tested the interactions again. Our hypothesis was that *Tc*MRGx/*Tc*MRGBP interaction would be less affected by changing R138 than *Tc*BDF5-MRG/*Tc*BDF5BP by changing R623. We explored this intervention by Y2H, and the results are shown in the **Figure *6*b**. As predicted, the interaction was completely abolished for *Tc*BDF5-MRG-R623A/*Tc*BDF5BP, while *Tc*MRG-R138A/*Tc*MRGBP interaction was almost unchanged. This suggests that the [FY]EPP motif is essential to maintain interaction by contacting the R623 of *Tc*BDF5.

In general, this shows again the level of detail reached by the models produced by AF2-multimer in trypanosomatids, since we can perform predictions with them, allowing us to get information about the importance of conserved sequences in interactions.

## Discussion

In this work, we addressed the problem of structurally defining the chromatin remodeling complexes from trypanosomatids. We identify two heterotrimers that assemble through interactions with MRG domains: One partially conserved, *Tc*TINTIN; and another completely novel, the one formed by *Tc*BDF5. We achieve this using several bioinformatics methods and subsequent validation by yeast hybrid approaches. We took advantage of having the AF2 monomer database available, as well as AF2-multimer available through ColabFold. This allows us to overcome limitations due to very low similarity (less than 30%) and original domain architecture, as found in proteins from deep branched protists, like trypanosomatids. For example, *Tc*MRGBP has a 17% sequence identity with its ortholog *Hs*MRGBP, making its identification impossible using highly performant algorithms such as HHblits^30^. However, AF2 was able to model the conserved fold of its globular domain, helping us in the identification. In this case, we approached the problem using the expert’s eye, since we extensively reviewed the literature describing the structures of complexes formed through MRG domains. Therefore, after reducing the number of potential MRGBPs by sequence analysis, we identified immediately that we were in presence of an MRGBP while reviewing the AF2-multimer predicted structures. In this sense, it is reasonable to think about the use of AF2 predicted structures to annotate sequences by structural similarity, automating the process of far homology detection. For example, Foldseek^48^ enables fast and sensitive comparisons of large structure sets and has a public server that finds structural similarities between any input PDB and the AF2 database.

On the other hand, the sequential assembly capacity found for *Tc*TINTIN may be associated with regulatory events that limit the formation of this complex to specific situations or locations where it is necessary. Another potential function could be to regulate the binding of BDF6 to acetyl-K. Given the BC loop of BD6 is involved in the interaction with *Tc*MRGBP, it is possible that the activation of this interaction by *Tc*MRGx regulates the acetyl-lysine(s) recognition capability of the hydrophobic pocket of BD6, either by enhancing its release from acetylated nucleosomes or by promoting its binding. For example, in humans, MRG15 mediates the activation of ASH1L histone methyltransferase by releasing an autoinhibitory loop upon interaction^22^. Whatever the case, it is remarkable that AF2 predicts interactions that could only happen under certain circumstances. Complementing the predicted complexes with experimental evidence and molecular dynamics simulations, as we did with *Tc*TINTIN, can explore crucial aspects of protein-protein interactions that static AF2-multimer models do not consider.

On the other hand, the interaction modes of the structures predicted for the heterotrimer assembled through *Tc*BDF5-MRG are totally novel. The way in which the IDR of *Tc*BDF5BP folds into an extended-turn-extended conformation has not been described before. Even the MRG is quite divergent, possessing several large-disordered loops, making it bigger than usual (**Figure S8**). Also, the cavity where the Fx[LIV]P motif of *Tc*BDF8 binds had never been described for an MRGBP. The latter may be a unique trypanosomatid feature, due to the divergence of this complex, or a yet undescribed interaction way present in eukaryotes. Nonetheless, *Tc*BDF8 engagement match the description of a bipartite interaction, present in all described MRGBPs^46^, as the IDR and DUF domains of *Tc*BDF8 were not able to express the reporter gene when separated. In this sense, we think this observation comes from the fact that both parts work together to bind *Tc*BDF5-MRG and stabilize the complex. One question that remains is if this comes from divergent or convergent evolution. Given these observations, is possible that a duplication of an MRG domain containing gene occurred at some point in evolution, opening the way to new epigenetic regulation mechanisms through genetic drift. Then, the MRG domain of BDF5 would be the consequence of said duplication and traces of this event could be found in other organisms. Supporting this, we identified an MRG-like fold in KAT2A residues 227-370 (**Figure S12**), one of the best-studied proteins with HAT activity in the human genome^49^, never reported to have an MRG-like structure. Far from stablishing that this domain is the ortholog of the MRG domain of BDF5, observations like this point to a picture more complicated than what it seems, and AF2 proves to be a great tool to clear it up.

We also explored the implication of the conserved arginines of *Tc*BDF5 and *Tc*MRGx in maintaining the interaction with the IDRs of *Tc*BDF5BP and *Tc*MRGBP, respectively. We verified through mutagenesis assays that the union between the aromatic ring of *Tc*BDF5BP-Y28 and the aliphatic chain of *Tc*BDF5-R623 is essential to sustain the union. Given the essentiality of BDF5^18^, an inhibitor could be developed that binds to the cavity formed above *Tc*BDF5-R623, where *Tc*BDF5BP-Y28 deepens, which is relatively different from that of *Hs*MRG15 and *Hs*MRGx. Alternatively, a PROTAC structure can be synthetized by, for example, linking the YEPP peptide to known E3 ligase ligands and assess whether this molecule manages to direct *Tc*BDF5 to its degradation^50^. Furthermore, with R623A mutant, we now have a model to study the biological implications of the *Tc*BDF5/*Tc*BDF5BP interaction and investigate its essentiality.

Finally, during the search, the structural models of many other protein monomers and complexes present in the datasets were also analyzed. Some of them could be in charge of assembling the even bigger complexes of which TINTIN or BDF5 are a part, but exploring this idea was beyond the objective of this work. However, we realize that with enough computing power, it is possible to compute the structure of the hypothetical megadalton complex of the BDF5-HAT2 network (CRKT), as well as the structure of the entire HAT1-containing NuA4 complex. For that, AF2Complex could be used, designed to predict the structure of large multimeric complexes using high-performance computing (HPC)^51^. Despite the implementation to be used, we proved that AF2 algorithm can be used to decipher the structure of complexes formed by very low, or even undetectable, sequence identity protein components, using interactomic datasets as starting search database.

## Methods

### Ortholog identification, alignments, and disorder prediction

The identification of orthologous proteins was performed using the OrthoMCL^52^ and TriTrypDB^35^ databases, with the IDs of the sequences present in the *Tb*HAT1 and *Tb*BDF5 interactomes as search input (Table 1 and Table 2, respectively), filtering by organism in the cases in which it was required. Multiple sequence alignment was carried out using ClustalO with default parameters. Disorder prediction was done with the software PrDOS^37^, using as query the sequences identified as orthologs from *T. cruzi* CL Brener strain.

### Multiple sequence alignment, structure prediction, visualization, and comparison

Multiple sequence alignments visualization was carried out using Jalview^53^. The monomer structures for each *T. cruzi* CL Brener ortholog were downloaded from AF2 protein structure database^39^. To map the predicted disorder and the degree of conservation into the structures, we setup Jalview to display them using PyMOL^54^, representing these annotations as color codes. The conservation annotation was performed by setting the threshold for color display using a sequence conservation degree of at least 30%.

Structure predictions of the complexes were preformed using ColabFold-MMseqs2 notebook^55^. `msa_mode` was set to `MMseqs2(UniRef+Environmental)`, `pair_mode` to `unpaired+paired, model_type` to `AlphaFold2-multimer-v2` and `num_recycles` to 3.

Structure alignment and comparison was performed in ChimeraX using matchmaker algorithm, guided by Smith and Waterman algorithm, which also provides the final RMSD values.

### CDS, cloning and expression vectors

All the cloning experiments were made using the Gateway cloning system^56^. Although the structures were predicted using the reference strain, the CDSs were cloned from the genomic DNA of Dm28c strain. Sequences ID and primers are shown in Table S1. The sequences were amplified by PCR, purified, and digested with the corresponding restriction enzymes, ligated into pENTR3C, transformed into electrocompetent DH5α cells and sequenced. The CDSs of *Tc*BDF5-MRG and *Tc*MRGBP were transferred to the Y2H vector pGBKT7-GW, leaving them in frame with the BD domain of GAL4 and a Myc-tag. The rest of the CDSs were transferred to the Y2H vector pGADT7-GW, leaving them in frame with the AD domain of GAL4 and an HA-tag. To perform the Y3H, we converted the yeast expression plasmid p426GPD to the gateway system, inserting the Gateway Cassette frame B (Invitrogen) into its SmaI site. To track whether *Tc*MRGx and *T*cBDF5-MRG were present in the nucleus, we N-terminal inserted the CDS of GFP in frame with both proteins into the vectors pENTR3C_*Tc*MRGx and pENTR3C_*Tc*BDF5-MRG. Finally, we transferred GFP, GFP-*Tc*MRGx and GFP-*Tc*BDF5-MRG constructs from the entry vectors to pGPD426-GW. The R to A mutations of the MRG domains were made by site directed mutagenesis of whole plasmids ^57^ on pENTR3C and after that transferred by recombination to pGBKT7-GW. All sequences were verified in the resulting plasmids by sequencing.

### Yeast strains

For the Y2H and Y3H experiments, *S. cerevisiae* MaV203 strain was used (*MAT*α, *leu*2-3,112, *trp*1-901, *his*3Δ200, *ade*2-101, *gal*4Δ, *gal*80Δ, *SPAL*10::URA3, *GAL*1::*lac*Z, *HIS*3_UAS GAL1_::*HIS*3@*LYS*2, *can*1^R^, *cyh*2^R^). For transformations, electrocompetent cells were prepared from a 1/10,000 dilution of a saturated culture that was inoculated into 500 ml liquid YPAD medium at 30°C and grown ON to OD_600_=1,0. Cells were cooled and centrifuged at 4000 g for 5 min. The supernatant was discarded, and the cells were washed twice with sterile distilled water and twice with 10% sorbitol. Cells were resuspended in 2 ml of 10% sorbitol and 40 μl per transformation mixed with up to 5 μl of plasmids were electroporated in 2 mm cuvettes (Gene Pulser Xcell BIO-RAD). For autoactivation controls, only one of the plasmids containing protein fusions was mixed with either empty pGBKT7 or empty pGADT7 (*e*.*g*., pGBKT7/pGADT7-GW_*Tc*MRGx). For the interaction tests cells, a combination of two plasmids containing two fusion proteins was used (*e*.*g*., pGBKT7-GW_*Tc*BDF6/pGADT7-GW_*Tc*MRGx). Electroporated cells were recovered in 1 ml of liquid YPAD with 10% sorbitol for 1 h at 30°C - 200 rpm and plated in Synthetic Complete (SC) medium lacking Trp to select pGBKT7 and lacking Leu for pGADT7 (SC-Leu-Trp).

The strains used in the Y3H experiments of *Tc*TINTIN were derived from the parental strains MaV203 pGADT7-GW_*Tc*MRGBP pGBKT7-GW_*Tc*BDF6 (interaction test), MaV203 pGADT7-GW_*Tc*MRGBP pGBKT7 and MaV203 pGADT7 pGBKT7-GW_*Tc*BDF6 (autoactivation controls), by introducing either p426GPD-GW_GFP or p426GPD-GW_GFP-*Tc*MRGx. They were generated using protocols mentioned above. Electroporated cells were selected using SC-Leu-Trp-Ura, to select for p426GPD-GW by also removing Ura. Only fluorescent colonies were used in the interaction assays. For the Y3H experiment of the heterotrimer formed by *Tc*BDF5-MRG the following cell strains were generated: MaV203 pGADT7-GW_*Tc*MRGBP pGBKT7-GW_*Tc*BDF8 p426GPD-GW_GFP (GFP control), MaV203 pGADT7 pGBKT7-GW_*Tc*BDF8 p426GPD-GW_GFP-*Tc*BDF5-MRG (autoactivation control 1), MaV203 pGADT7-GW_*Tc*MRGBP pGBKT7 p426GPD-GW_GFP-*Tc*BDF5-MRG (autoactivation control 2) and MaV203 pGADT7-GW_*Tc*MRGBP pGBKT7-GW_*Tc*BDF8 p426GPD-GW_GFP-*Tc*BDF5-MRG (interaction test).

### Y2H and Y3H assays

**Master plating**: For each experiment, desired colonies were first streaked onto master plates with SC-Leu-Trp for Y2H experiments or SC-Leu-Trp-Ura for Y3H and grown for 24 h at 30°C. Four different clones were streaked for the interaction tests and two for each of the controls. **Replica plating:** The master plates were replicated using autoclaved velvets, transferring the streaks onto a nitrocellulose membrane supported on a YPAD plate to evaluate the expression of β-galactosidase. **X-Gal assay:** After incubation, nitrocellulose membranes were removed from YPAD plates and immersed in liquid N_2_ for 20-30 s. Membranes were then transferred on top of round Whatman filters soaked with X-gal buffer (10 mg X-Gal, 100 μl DMF, 10 ml Z-buffer) colonies side up. The appearance of blue color was monitored over a 24 h period at 37°C. **Spot growth assay**. Spot growth assays were conducted to use *HIS3* as reporter gene. MaV203 cells transformed with plasmids derived from pGBKT7-GW and pGADT7-GW were grown overnight (ON) in liquid SC-Leu-Trp medium at 30°C with vigorous shaking. Yeast cells also transformed with plasmids derived from p426GPD-GW were grown ON in liquid SC-Leu-Trp-Ura medium. The cells were suspended at OD_600_=1,000 and serially diluted by 1/10. 7 μl of each culture dilution was spotted on solid SC medium alone or solid SC-His supplemented with 50 mM of 3-amino-1,2,4-triazole (3-AT) to titrate basal *HIS3* expression. Plates were incubated at 30°C and followed for 4 to 12 days.

### Western blot

Cells were inoculated and grown for 48 h at 200 rpm and 30°C in 5 ml of the adequate liquid SC medium required to keep selective pressure. Cells were harvested by centrifugation (8000 g, 5 min) and the pellet was resuspended in 200 μl of extraction buffer (0.1 M NaOH, 0.05 M EDTA, 2% SDS, 2% β-mercaptoethanol). After 10 min at 96°C, 5 μl of 4 M acetic acid was added, followed by 50 μl of loading buffer. 8 μL of each sample was loaded per lane onto a 15% SDS-PAGE gel. After electrophoresis, the proteins were transferred to a nitrocellulose membrane and stained with Ponceau Red stain to ensure even loading and proper transfer. Blocking was done ON with PBS - 0.1% Tween20 (PBST) - 5% non-fat milk at 4°C. 3.3 μg/ml of rabbit anti-Myc primary antibody (Calbiochem) was incubated for 1 h at room temperature (RT) in PBST 5% milk. Three 10 min washes were performed with 5 ml of PBST. Goat secondary antibody anti-rabbit (GE) conjugated to HRP was incubated for 1 hour at RT (1/20000 dilution) in PBST 5% milk. The membrane was then stripped and incubated with rat anti-HA primary antibody (Roche) at 1/2000 dilution in PBST 5% milk and washed 3 times. Goat secondary antibody anti-rat (Pierce) conjugated to HRP was incubated for 1 hs at RT (1/10000 dilution) in PBST 5% milk. For the detection of GFP tagged proteins, mouse anti-GFP primary antibody (GE) was used (1/2000 dilution). The incubation was performed for 3 hs at RT in PBST – 5% non-fat milk. The membrane was then washed and incubated 1 hs at RT with goat anti-mouse secondary antibody conjugated to HRP (Roche) at 1/20000 dilution in PBST - 5% milk.

### Phylogenetic analysis

The FHA protein sequences for constructing the phylogenetic tree were retrieved from Pfam^43^ entry PF00498, restricting the download only to seed sequences and requesting them pre-aligned. Residues 310-472 from TcCLB.510747.120, 328-507 from Tb927.9.13320 and 342-555 from LINF_350030200, corresponding to the FHA domains, were added to the multiple sequence alignment using MAFFT^58^ version 7 setting the option --`add` to maintain the original alignment. Positions with low phylogenetic information were filtered using ClipKIT^59^ using the trimming mode -`m kpic-gappy`. The phylogenetic tree was inferred by maximum-likelihood using IQ-TREE^60^, setting a bootstrap of 1000 and using ModelFinder^61^ to find the best substitution model. Tree visualizations were made with Dendroscope^62^.

### Molecular dynamics simulations and analysis

*Tc*MRGx/*Tc*MRGBP dimer and *Tc*MRGBP monomer structures were protonated according to the pKa calculated by PROPKA v3^63^. Systems involving both models were then prepared in CHARMM-GUI^64^. A solvation box of 15Å thickness was added to each model using the TIP3 model. Then the systems were neutralized using 0.15M of NaCl. After assembling the two systems, we performed an optimization using the steep descent method with 50000 steps. Then, we performed heating of the systems from 100K to 310K in ensemble NVT, during 7.5ns. Next, we performed an equilibration step of the systems, in ensemble NPT (310K and 1atm) by 7.5ns using v-rescale thermostat and c-rescale barostat. After the equilibration step, we perform the production step for 200ns in ensemble NPT using the leapfrog algorithm with an integration step of 2fs. For temperature and pressure control, we used the Nose-Hoover and Parrinello-Rahman algorithms, respectively. We also used the LINCS algorithm to control the H-bonds. For short-range electrostatic interactions and Van der Waals interactions, we apply a cut-off of 10Å. The algorithm for electrostatic interactions integration applied by us was PME. The force field used in the simulations was CHARMM36^65^. For both systems, we performed three replicas changing the seed of the atom’s initial velocity. All molecular dynamics simulations were performed with GROMACS v2020.3^66^.

To perform a structural analysis, conformations from MD simulations were saved each 20ps. RMSD, RMSF, radius of gyration (RoG), and SASA values were calculated for each set of structures using pytraj and mdtraj python packages. To obtain clusters and medoid structures, we concatenated the three replicas for each condition and an allxall RMSD matrix considered as reaction coordinate. The PCA method was used to reduce the data dimensionality and clustering was performed using the agglomerative ward algorithm. We used the WHAM method to obtain a free-energy graph of structures sampled from MD simulations. We used one-way ANOVA with F-test to verify the significance in RMSD, RoG, and SASA average variation.

## Data availability

Complex structure predictions have been uploaded as comments into the corresponding gene entries from TriTrypDB *T. cruzi* CL Brener strain. *Tc*TINTIN complexes: TcCLB.505999.50 (*Tc*MRGx), TcCLB.511217.190 (*Tc*MRGBP) and TcCLB.506529.660 (*Tc*BDF6). *Tc*BDF5-MRG complexes: TcCLB.506755.120 (*Tc*BDF5), TcCLB.510747.120 (*Tc*BDF5BP) and TcCLB.506559.310 (*Tc*BDF8).

## Acknowledgments

This work was funded by the National Research Council, CONICET (PIP 2021-0848), the National Agency for Science, ANPCyT (PICT 2017-1978, 2020-01704) and Universidad Nacional de Rosario (0001-00285485). We want to thank Dr. Antonio Uttaro, who kindly give us the p426GPD plasmid.

## References

(1) Gazestani, V. H.; Yip, C. W.; Nikpour, N.; Berghuis, N.; Salavati, R. TrypsNetDB: An Integrated Framework for the Functional Characterization of Trypanosomatid Proteins. PLoS Negl Trop Dis 2017, 11 (2), 1–11. https://doi.org/10.1371/journal.pntd.0005368.

(2) Büscher, P.; Cecchi, G.; Jamonneau, V.; Priotto, G. Human African Trypanosomiasis. The Lancet 2017, 390 (10110), 2397–2409. https://doi.org/10.1016/S0140-6736(17)31510-6.

(3) Pérez-Molina, J. A.; Molina, I. Chagas Disease. The Lancet 2018, 391 (10115), 82–94. https://doi.org/10.1016/S0140-6736(17)31612-4.

(4) Burza, S.; Croft, S. L.; Boelaert, M. Leishmaniasis. The Lancet 2018, 392 (10151), 951–970. https://doi.org/10.1016/S0140-6736(18)31204-2.

(5) De Gaudenzi, J. G.; Noé, G.; Campo, V. A.; Frasch, A. C.; Cassola, A. Gene Expression Regulation in Trypanosomatids. Essays Biochem 2011, 51, 31–46. https://doi.org/10.1042/bse0510031.

(6) Gilinger, G.; Bellofatto, V. Trypanosome Spliced Leader RNA Genes Contain the First Identified RNA Polymerase II Gene Promoter in These Organisms. Nucleic Acids Res 2001, 29 (7), 1556–1564. https://doi.org/10.1093/nar/29.7.1556.

(7) Cordon-Obras, C.; Gomez-Liñan, C.; Torres-Rusillo, S.; Vidal-Cobo, I.; Lopez-Farfan, D.; Barroso-del Jesus, A.; Rojas-Barros, D.; Carrington, M.; Navarro, M. Identification of Sequence-Specific Promoters Driving Polycistronic Transcription Initiation by RNA Polymerase II in Trypanosomes. Cell Rep 2022, 38 (2). https://doi.org/10.1016/j.celrep.2021.110221.

(8) Schimanski, B.; Laufer, G.; Gontcharova, L.; Günzl, A. The Trypanosoma Brucei Spliced Leader RNA and RRNA Gene Promoters Have Interchangeable TbSNAP50-Binding Elements. Nucleic Acids Res 2004, 32 (2), 700–709. https://doi.org/10.1093/nar/gkh231.

(9) Saha, S. Histone Modifications and Other Facets of Epigenetic Regulation in Trypanosomatids: Leaving Their Mark. mBio 2020, 11 (5), 1–19. https://doi.org/10.1128/mBio.01079-20.

(10) Kouzarides, T. Chromatin Modifications and Their Function. Cell 2007, 128 (4), 693–705. https://doi.org/10.1016/j.cell.2007.02.005.

(11) Biswas, S.; Rao, C. M. Epigenetic Tools (The Writers, The Readers and The Erasers) and Their Implications in Cancer Therapy. Eur J Pharmacol 2018, 837 (August), 8–24. https://doi.org/10.1016/j.ejphar.2018.08.021.

(12) Yang, X. J.; Seto, E. HATs and HDACs: From Structure, Function and Regulation to Novel Strategies for Therapy and Prevention. Oncogene 2007, 26 (37), 5310–5318. https://doi.org/10.1038/sj.onc.1210599.

(13) Doyon, Y.; Côté, J. The Highly Conserved and Multifunctional NuA4 HAT Complex. Curr Opin Genet Dev 2004, 14 (2), 147–154. https://doi.org/10.1016/j.gde.2004.02.009.

(14) Joshi, P.; Greco, T. M.; Guise, A. J.; Luo, Y.; Yu, F.; Nesvizhskii, A. I.; Cristea, I. M. The Functional Interactome Landscape of the Human Histone Deacetylase Family. Mol Syst Biol 2013, 9 (1), 1–21. https://doi.org/10.1038/msb.2013.26.

(15) Ye, F.; Huang, J.; Wang, H.; Luo, C.; Zhao, K. Targeting Epigenetic Machinery: Emerging Novel Allosteric Inhibitors. Pharmacol Ther 2019, 204, 107406. https://doi.org/10.1016/j.pharmthera.2019.107406.

(16) Pezza, A.; Tavernelli, L. E.; Alonso, V. L.; Perdomo, V.; Gabarro, R.; Prinjha, R.; Rodríguez Araya, E.; Rioja, I.; Docampo, R.; Calderón, F.; Martin, J.; Serra, E. Essential Bromodomain TcBDF2 as a Drug Target against Chagas Disease. ACS Infect Dis 2022. https://doi.org/10.1021/acsinfecdis.2c00057.

(17) Staneva, D. P.; Carloni, R.; Auchynnikava, T.; Tong, P.; Rappsilber, J.; Jeyaprakash, A. A.; Matthews, K. R.; Allshire, R. C. A Systematic Analysis of Trypanosoma Brucei Chromatin Factors Identifies Novel Protein Interaction Networks Associated with Sites of Transcription Initiation and Termination. Genome Res 2021, 31 (11), 2138–2154. https://doi.org/10.1101/gr.275368.121.

(18) Jones, N. G.; Geoghegan, V.; Moore, G.; Carnielli, J. B. T.; Newling, K.; Calderón, F.; Gabarró, R.; Martín, J.; Prinjha, R. K.; Rioja, I.; Wilkinson, A. J.; Mottram, J. C. Bromodomain Factor 5 Is an Essential Regulator of Transcription in Leishmania. Nat Commun 2022, 13 (1), 4071. https://doi.org/10.1038/s41467-022-31742-1.

(19) Vellmer, T.; Hartleb, L.; Sola, A. F.; Kramer, S.; Meyer-Natus, E.; Butter, F.; Janzen, C. J. A Novel SNF2 ATPase Complex in Trypanosoma Brucei with a Role in H2A.Z-Mediated Chromatin Remodelling. PLoS Pathog 2022, 18 (6), 1–29. https://doi.org/10.1371/journal.ppat.1010514.

(20) Bowman, B. R.; Moure, C. M.; Kirtane, B. M.; Welschhans, R. L.; Tominaga, K.; Pereira-Smith, O. M.; Quiocho, F. A. Multipurpose MRG Domain Involved in Cell Senescence and Proliferation Exhibits Structural Homology to a DNA-Interacting Domain. Structure 2006, 14 (1), 151–158. https://doi.org/10.1016/j.str.2005.08.019.

(21) Xie, T.; Zmyslowski, A. M.; Zhang, Y.; Radhakrishnan, I. Structural Basis for Multi-Specificity of MRG Domains. Structure 2015, 23 (6), 1049–1057. https://doi.org/10.1016/j.str.2015.03.020.

(22) Lee, Y.; Yoon, E.; Cho, S.; Schmähling, S.; Müller, J.; Song, J. J. Structural Basis of MRG15-Mediated Activation of the ASH1L Histone Methyltransferase by Releasing an Autoinhibitory Loop. Structure 2019, 27 (5), 846-852.e3. https://doi.org/10.1016/j.str.2019.01.016.

(23) Sapoval, N.; Aghazadeh, A.; Nute, M. G.; Antunes, D. A.; Balaji, A.; Baraniuk, R.; Barberan, C. J.; Dannenfelser, R.; Dun, C.; Edrisi, M.; Elworth, R. A. L.; Kille, B.; Kyrillidis, A.; Nakhleh, L.; Wolfe, C. R.; Yan, Z.; Yao, V.; Treangen, T. J. Current Progress and Open Challenges for Applying Deep Learning across the Biosciences. Nat Commun 2022, 13 (1), 1–12. https://doi.org/10.1038/s41467-022-29268-7.

(24) Jumper, J.; Evans, R.; Pritzel, A.; Green, T.; Figurnov, M.; Ronneberger, O.; Tunyasuvunakool, K.; Bates, R.; Žídek, A.; Potapenko, A.; Bridgland, A.; Meyer, C.; Kohl, A. A.; Ballard, A. J.; Cowie, A.; Romera-Paredes, B.; Nikolov, S.; Jain, R.; Adler, J.; Back, T. ; Petersen, S.; Reiman, D.; Clancy, E.; Zielinski, M.; Steinegger, M.; Pacholska, M.; Berghammer, T.; Bodenstein, S.; Silver, D.; Vinyals, O.; Senior, A. W.; Kavukcuoglu, K.; Kohli, P.; Hassabis, D. Highly Accurate Protein Structure Prediction with AlphaFold. Nature 2021, 596 (7873), 583–589. https://doi.org/10.1038/s41586-021-03819-2.

(25) Evans, R.; O’Neill1, M.; Pritzel1, A.; Antropova, N.; Senior, A.; Green, T.; Žídek, A.; Bates, R.; Blackwell, S.; Yim, J.; Ronneberger, O.; Bodenstein, S.; Zielinski, M.; Bridgland, A.; Potapenk, A.; Clancy, E.; Kohli, P.; Jumper, J.; Hassabis, D. Protein Complex Prediction with AlphaFold2-Multimer. Methods in Molecular Biology 2021, 804, 297–312. https://doi.org/doi.org/10.1101/2021.10.04.463034.

(26) Bhat, W.; Ahmad, S.; Côté, J. TINTIN, at the Interface of Chromatin, Transcription Elongation, and MRNA Processing. RNA Biol 2015, 12 (5), 486–489. https://doi.org/10.1080/15476286.2015.1026032.

(27) Devoucoux, M.; Roques, C.; Lachance, C.; Lashgari, A.; Joly-Beauparlant, C.; Jacquet, K.; Alerasool, N.; Prudente, A.; Taipale, M.; Droit, A.; Lambert, J.-P.; Hussein, S. M. I.; Côté, J. MRG Proteins Are Shared by Multiple Protein Complexes with Distinct Functions. Molecular & Cellular Proteomics 2022, 21, 100253. https://doi.org/10.1016/j.mcpro.2022.100253.

(28) Jones, N. G.; Geoghegan, V.; Moore, G.; Carnielli, J. B. T.; Newling, K.; Calderón, F.; Gabarró, R.; Martín, J.; Prinjha, R. K.; Rioja, I.; Wilkinson, A. J.; Mottram, J. C. Bromodomain Factor 5 Is an Essential Regulator of Transcription in Leishmania. Nat Commun 2022, 13 (1), 4071. https://doi.org/10.1038/s41467-022-31742-1.

(29) Staneva, D. P.; Carloni, R.; Auchynnikava, T.; Tong, P.; Arulanandam, J. A.; Rappsilber, J.; Matthews, K. R.; Allshire, R. A Systematic Analysis of Trypanosoma Brucei Chromatin Factors Identifies Novel Protein Interaction Networks Associated with Sites of Transcription Initiation and Termination. bioRxiv 2021, 2021.02.09.430399.

(30) Remmert, M.; Biegert, A.; Hauser, A.; Söding, J. HHblits: Lightning-Fast Iterative Protein Sequence Searching by HMM-HMM Alignment. Nat Methods 2012, 9 (2), 173–175. https://doi.org/10.1038/nmeth.1818.

(31) Cheung, A. C. M.; Díaz-Santín, L. M. Share and Share Alike: The Role of Tra1 from the SAGA and NuA4 Coactivator Complexes. Transcription 2019, 10 (1), 37–43. https://doi.org/10.1080/21541264.2018.1530936.

(32) Kraus, A. J.; Vanselow, J. T.; Lamer, S.; Brink, B. G.; Schlosser, A.; Siegel, T. N. Distinct Roles for H4 and H2A.Z Acetylation in RNA Transcription in African Trypanosomes. Nat Commun 2020, 11 (1). https://doi.org/10.1038/s41467-020-15274-0.

(33) Almawi, A. W.; Matthews, L. A.; Guarné, A. FHA Domains: Phosphopeptide Binding and Beyond. Prog Biophys Mol Biol 2017, 127, 105–110. https://doi.org/10.1016/j.pbiomolbio.2016.12.003.

(34) Cribb, P.; Serra, E. One- and Two-Hybrid Analysis of the Interactions between Components of the Trypanosoma Cruzi Spliced Leader RNA Gene Promoter Binding Complex. Int J Parasitol 2009, 39 (5), 525–532. https://doi.org/10.1016/j.ijpara.2008.09.008.

(35) Amos, B.; Aurrecoechea, C.; Barba, M.; Barreto, A.; Basenko, E. Y.; Ba, W.; Belnap, R.; Blevins, A. S.; Ulrike, B.; Brestelli, J.; Brunk, B. P.; Caddick, M.; Callan, D.; Campbell, L.; Christensen, M. B.; Christophides, G. K.; Crouch, K.; Davis, K.; Debarry, J.; Doherty, R.; Duan, Y.; Dunn, M.; Falke, D.; Fisher, S.; Flicek, P.; Fox, B.; Gajria, B.; Giraldo-calder, G. I.; Harb, O. S.; Harper, E.; Hertz-fowler, C.; Hickman, M. J.; Howington, C.; Hu, S.; Humphrey, J.; Iodice, J.; Jones, A.; Judkins, J.; Kelly, S. A.; Kissinger, J. C.; Kwon, D. K.; Lamoureux, K.; Lawson, D.; Li, W.; Lies, K.; Lodha, D.; Long, J.; Maccallum, R. M.; Maslen, G.; Mcdowell, M. A.; Nabrzyski, J.; Roos, D. S.; Rund, S. S. C.; Schulman, S. W.; Shanmugasundram, A.; Sitnik, V.; Spruill, D.; Starns, D.; Stoeckert, C. J.; Tomko, S. S.; Wang, H.; Warrenfeltz, S.; Wieck, R.; Wilkinson, P. A.; Xu, L.; Zheng, J. VEuPathDB : The Eukaryotic Pathogen, Vector and Host Bioinformatics Resource Center. 2021, 1–14.

(36) Sievers, F.; Higgins, D. G. Clustal Omega for Making Accurate Alignments of Many Protein Sequences. Protein Science 2018, 27 (1), 135–145. https://doi.org/10.1002/pro.3290.

(37) Ishida, T.; Kinoshita, K. PrDOS: Prediction of Disordered Protein Regions from Amino Acid Sequence. Nucleic Acids Res 2007, 35 (SUPPL.2), 460–464. https://doi.org/10.1093/nar/gkm363.

(38) Wilson, C. J.; Choy, W. Y.; Karttunen, M. AlphaFold2: A Role for Disordered Protein/Region Prediction? Int J Mol Sci 2022, 23 (9), 1–14. https://doi.org/10.3390/ijms23094591.

(39) Varadi, M.; Anyango, S.; Deshpande, M.; Nair, S.; Natassia, C.; Yordanova, G.; Yuan, D.; Stroe, O.; Wood, G.; Laydon, A.; Zídek, A.; Green, T.; Tunyasuvunakool, K.; Petersen, S.; Jumper, J.; Clancy, E.; Green, R.; Vora, A.; Lutfi, M.; Figurnov, M.; Cowie, A.; Hobbs, N.; Kohli, P.; Kleywegt, G.; Birney, E.; Hassabis, D.; Velankar, S. AlphaFold Protein Structure Database: Massively Expanding the Structural Coverage of Protein-Sequence Space with High-Accuracy Models. Nucleic Acids Res 2022, 50 (D1), D439–D444. https://doi.org/10.1093/nar/gkab1061.

(40) Wang, X.; Ahmad, S.; Zhang, Z.; Côté, J.; Cai, G. Architecture of the Saccharomyces Cerevisiae NuA4/TIP60 Complex. Nat Commun 2018, 9 (1), 1–11. https://doi.org/10.1038/s41467-018-03504-5.

(41) Invitrogen. ProQuest TM Two-Hybrid System with Gateway ® Technology; 2002.

(42) Zhang, B. W.; Arasteh, S.; Levy, R. M. The UWHAM and SWHAM Software Package. Sci Rep 2019, 9 (1), 1–9. https://doi.org/10.1038/s41598-019-39420-x.

(43) Mistry, J.; Chuguransky, S.; Williams, L.; Qureshi, M.; Salazar, G. A.; Sonnhammer, E. L. L.; Tosatto, S. C. E.; Paladin, L.; Raj, S.; Richardson, L. J.; Finn, R. D.; Bateman, A. Pfam: The Protein Families Database in 2021. Nucleic Acids Res 2021, 49 (D1), D412–D419. https://doi.org/10.1093/nar/gkaa913.

(44) Cai, Y.; Jin, J.; Swanson, S. K.; Cole, M. D.; Choi, S. H.; Florens, L.; Washburn, M. P.; Conaway, J. W.; Conaway, R. C. Subunit Composition and Substrate Specificity of a MOF-Containing Histone Acetyltransferase Distinct from the Male-Specific Lethal (MSL) Complex. Journal of Biological Chemistry 2010, 285 (7), 4268–4272. https://doi.org/10.1074/jbc.C109.087981.

(45) Kadlec, J.; Hallacli, E.; Lipp, M.; Holz, H.; Sanchez-Weatherby, J.; Cusack, S.; Akhtar, A. Structural Basis for MOF and MSL3 Recruitment into the Dosage Compensation Complex by MSL1. Nat Struct Mol Biol 2011, 18 (2), 142–150. https://doi.org/10.1038/nsmb.1960.

(46) Redington, J.; Deveryshetty, J.; Kanikkannan, L.; Miller, I.; Korolev, S. Structural Insight into the Mechanism of Palb2 Interaction with Mrg15. Genes (Basel) 2021, 12 (12). https://doi.org/10.3390/genes12122002.

(47) Hou, P.; Huang, C.; Liu, C. P.; Yang, N.; Yu, T.; Yin, Y.; Zhu, B.; Xu, R. M. Structural Insights into Stimulation of Ash1L’s H3K36 Methyltransferase Activity through Mrg15 Binding. Structure 2019, 27 (5), 837-845.e3. https://doi.org/10.1016/j.str.2019.01.015.

(48) Kempen, M. van; Kim, S. S.; Tumescheit, C.; Mirdita, M.; Gilchrist, C. L. M.; Söding, J.; Steinegger, M. Foldseek: Fast and Accurate Protein Structure Search. bioRxiv 2022, 1, 2022.02.07.479398.

(49) Haque, Md. E.; Jakaria, Md.; Akther, M.; Cho, D.-Y.; Kim, I.-S.; Choi, D.-K. The GCN5: Its Biological Functions and Therapeutic Potentials. Clin Sci 2021, 135 (1), 231–257. https://doi.org/10.1042/CS20200986.

(50) Békés, M.; Langley, D. R.; Crews, C. M. PROTAC Targeted Protein Degraders: The Past Is Prologue. Nat Rev Drug Discov 2022, 21 (3), 181–200. https://doi.org/10.1038/s41573-021-00371-6.

(51) Gao, M.; Nakajima An, D.; Parks, J. M.; Skolnick, J. AF2Complex Predicts Direct Physical Interactions in Multimeric Proteins with Deep Learning. Nat Commun 2022, 13 (1), 1744. https://doi.org/10.1038/s41467-022-29394-2.

(52) Li, L.; Stoeckert, C. J. J.; Roos, D. S. OrthoMCL: Identification of Ortholog Groups for Eukaryotic Genomes. Genome Res 2003, 13 (9), 2178–2189. https://doi.org/10.1101/gr.1224503.

(53) Waterhouse, A. M.; Procter, J. B.; Martin, D. M. A.; Clamp, M.; Barton, G. J. Jalview Version 2-A Multiple Sequence Alignment Editor and Analysis Workbench. Bioinformatics 2009, 25 (9), 1189–1191. https://doi.org/10.1093/bioinformatics/btp033.

(54) Martinez, X.; Krone, M.; Alharbi, N.; Rose, A. S.; Laramee, R. S.; O’Donoghue, S.; Baaden, M.; Chavent, M. Molecular Graphics: Bridging Structural Biologists and Computer Scientists. Structure 2019, 27 (11), 1617–1623. https://doi.org/10.1016/j.str.2019.09.001.

(55) Mirdita, M.; Schütze, K.; Moriwaki, Y.; Heo, L.; Ovchinnikov, S.; Steinegger, M. ColabFold: Making Protein Folding Accessible to All. Nat Methods 2022, 19 (6), 679–682. https://doi.org/10.1038/s41592-022-01488-1.

(56) Reece-Hoyes, J. S.; Walhout, A. J. M. Gateway Recombinational Cloning. Cold Spring Harb Protoc 2018, 2018 (1), 1–6. https://doi.org/10.1101/pdb.top094912.

(57) Laible, M.; Boonrod, K. Homemade Site Directed Mutagenesis of Whole Plasmids. Journal of Visualized Experiments 2009, No. 27, 1–3. https://doi.org/10.3791/1135.

(58) Katoh, K.; Rozewicki, J.; Yamada, K. D. MAFFT Online Service: Multiple Sequence Alignment, Interactive Sequence Choice and Visualization. Brief Bioinform 2018, 20 (4), 1160–1166. https://doi.org/10.1093/bib/bbx108.

(59) Steenwyk, J. L.; Buida, T. J.; Li, Y.; Shen, X. X.; Rokas, A. ClipKIT: A Multiple Sequence Alignment Trimming Software for Accurate Phylogenomic Inference. PLoS Biol 2020, 18 (12), 1–17. https://doi.org/10.1371/journal.pbio.3001007.

(60) Minh, B. Q.; Schmidt, H. A.; Chernomor, O.; Schrempf, D.; Woodhams, M. D.; von Haeseler, A.; Lanfear, R.; Teeling, E. IQ-TREE 2: New Models and Efficient Methods for Phylogenetic Inference in the Genomic Era. Mol Biol Evol 2020, 37 (5), 1530–1534. https://doi.org/10.1093/molbev/msaa015.

(61) Kalyaanamoorthy, S.; Minh, B. Q.; Wong, T. K. F.; von Haeseler, A.; Jermiin, L. S. ModelFinder: Fast Model Selection for Accurate Phylogenetic Estimates. Nat Methods 2017, 14 (6), 587–589. https://doi.org/10.1038/nmeth.4285.

(62) Huson, D. H.; Scornavacca, C. Dendroscope 3: An Interactive Tool for Rooted Phylogenetic Trees and Networks. Syst Biol 2012, 61 (6), 1061–1067. https://doi.org/10.1093/sysbio/sys062.

(63) Olsson, M. H. M.; Søndergaard, C. R.; Rostkowski, M.; Jensen, J. H. PROPKA3: Consistent Treatment of Internal and Surface Residues in Empirical PKa Predictions. J Chem Theory Comput 2011, 7 (2), 525–537. https://doi.org/10.1021/ct100578z.

(64) Jo, S.; Kim, T.; Iyer, V. G.; Im, W. CHARMM-GUI: A Web-Based Graphical User Interface for CHARMM. J Comput Chem 2008, 29 (11), 1859–1865. https://doi.org/10.1002/jcc.20945.

(65) Huang, J.; MacKerell, A. D. J. CHARMM36 All-Atom Additive Protein Force Field: Validation Based on Comparison to NMR Data. J Comput Chem 2013, 34 (25), 2135–2145. https://doi.org/10.1002/jcc.23354.

(66) Berendsen, H. J. C.; van der Spoel, D.; van Drunen, R. GROMACS: A Message-Passing Parallel Molecular Dynamics Implementation. Comput Phys Commun 1995, 91 (1–3), 43–56. https://doi.org/10.1016/0010-4655(95)00042-e.

