## Supporting_Information for "Deciphering divergent trypanosomatid nuclear complexes by analyzing interactomic datasets with AlphaFold2 and genetic approaches"

#### Supporting figures

**Figure S1**

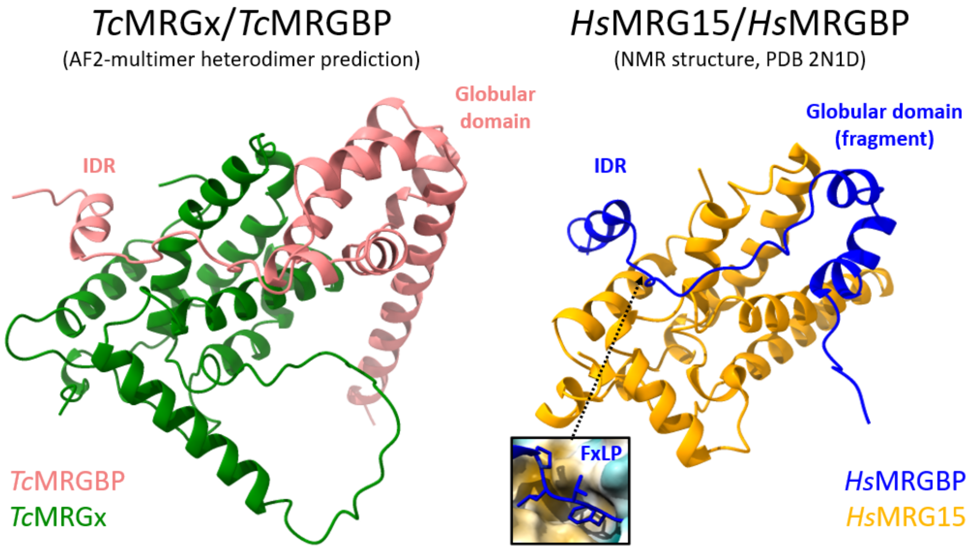


**Figure S1: Comparison between the predicted *Tc*MRGx/*Tc*MRGBP dimer and the solution structure 2N1D.** The best ranked model of the *Tc*MRGx/*Tc*MRGBP pair was aligned in ChimeraX using matchmaker algorithm with one of the conformers of the 2N1D solution structure and then displayed in two different panels. Inset shows the structure of the FxLP motif positioned on a pocket formed on the solvent-exposed surface of *Hs*MRG15 colored according to its degree of hydrophobicity (yellow = hydrophobic, light blue = hydrophilic). The arrow points to its position on the IDR of *Hs*MRGBP. The HsMRGBP sequence chosen to resolve by NMR in 2N1D omitted a fragment of the globular domain.

### Figure S2

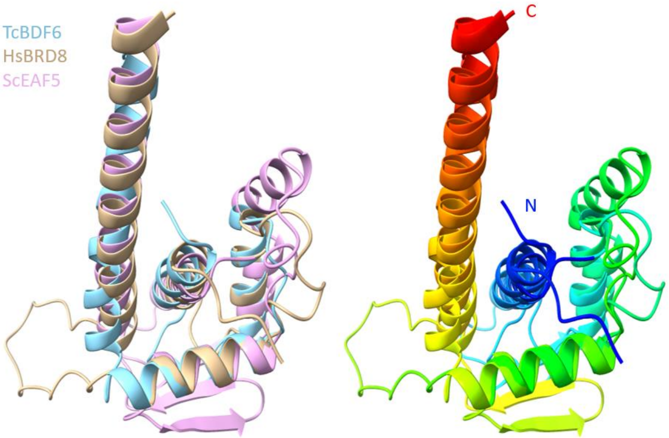


**Figure S2: The structure alignment of the N-terminal domains of *Tc*BDF6, *Hs*BRD8 and *Sc*EAF5 indicates that they are homologous.** On the left are the models of these domains colored my chain. On the right are the models colored in rainbow colors by position, from blue in the N-terminal residue to red in the C-terminal residue.

### Figure S3

| Protein 1 (DB fusion) | Protein 2 (AD fusion) |
|---|---|
| *Tc*MRGBP | *Tc*BDF6 |
| *Tc*MRGBP | *Tc*MRGx |
| *Tc*MRGBP | - |
| - | *Tc*BDF6 |
| - | *Tc*MRGx |
| Control (++) | |
| Control (+) | |
| Control (-) | |

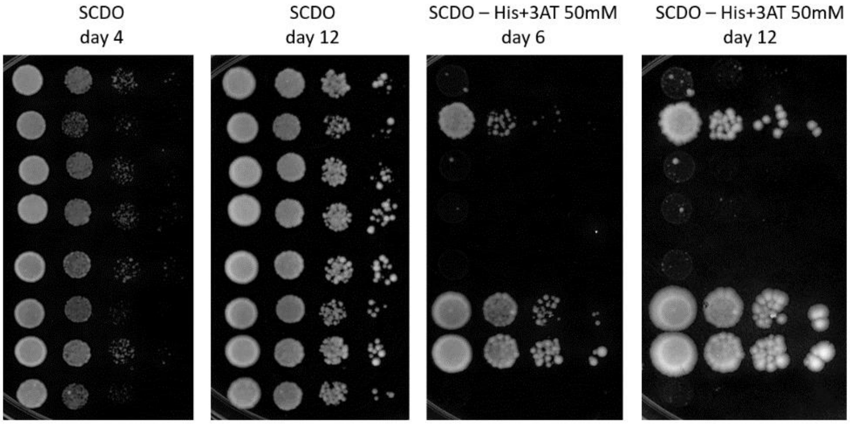


**Figure S3: Interaction tests by spot growth assay between *Tc*MRGBP and *Tc*BDF6 or *Tc*MRGx using HIS3 as reporter gene**. Spot growth assay of MaV203 strains expressing a combination of AD, DB, DB-*Tc*MRGBP, AD-*Tc*BDF6 and AD-*Tc*MRGx. The table on the left indicates the combinations tested in each row. Positive and negative interaction controls were also included. Dilutions of $OD_{600}$=1.000, 0.100, 0.010 and 0.001 were spotted on solid SC medium or solid SC medium without histidine (-His) supplemented with 50mM 3-AT to titrate basal HIS3 expression. Cells were incubated at 30 °C for 4 to 12 days.

### Figure S4

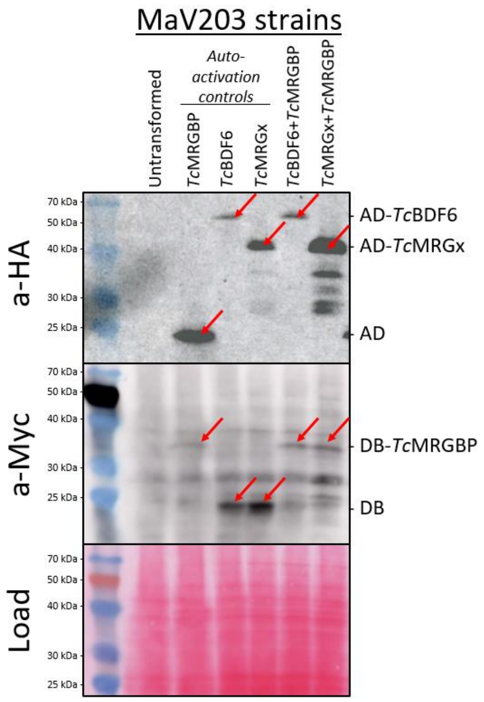


**Figure S4: Protein expression assessed by western blot.** Western blots of total extracts of the yeast strains from the master plate on **Error! Reference source not found.a**. The a-HA blot shows the bands corresponding to constructs containing the AD domain and the a-Myc blot shows the bands corresponding to constructs containing the DB domain. Ponceau stain (Load) displays even loading of the samples. Red arrows point to the bands corresponding to the expected sizes.

### Figure S5

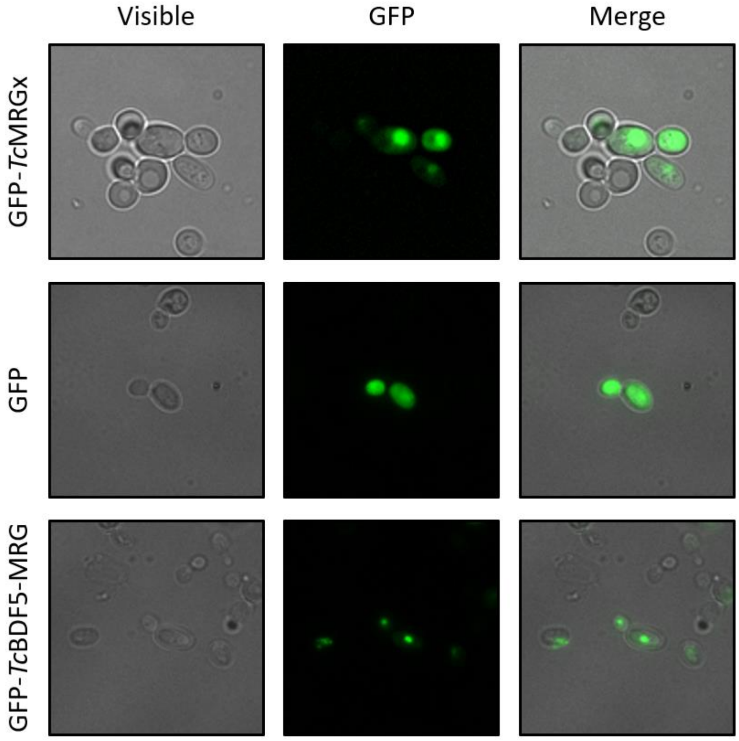


**Figure S5: Direct fluorescence of GFP-tagged proteins used in the Y3H experiments.** The first row shows nuclear green signal coming from GFP-*Tc*MRGx expression in the strain with the complete Y3H system from **Error! Reference source not found.c** (Full *Tc*TINTIN). The same signal was observed for the control strains expressing GFP-*Tc*MRGx from

**Figure S6**. The second row shows the signal of GFP alone distributed in all the compartments of the GFP control strain from **Figure S6**. The same pattern was seen for the GFP control on the Y3H experiment for *Tc*BDF5-MRG heterotrimer. The last row displays the green signal coming from GFP-*Tc*BDF5-MRG, expressed in the strain with the complete Y3H system from **Figure 5d**. The same pattern was observed for the rest of the control strains expressing GFP-*Tc*BDF5-MRG.

### Figure S6

| Protein 1 | Protein 2 | Protein 3 |
|---|---|---|
| DB-*Tc*MRGBP | GFP-*Tc*MRGx | AD-*Tc*BDF6 |
| DB-*Tc*MRGBP | GFP | AD-*Tc*BDF6 |
| DB | GFP-*Tc*MRGx | AD-*Tc*BDF6 |
| DB-*Tc*MRGBP | GFP-*Tc*MRGx | AD |
| | Control (++) | |
| | Control (-) | |

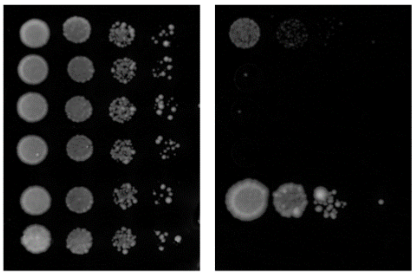


**Figure S6: *Tc*MRGx activates the interaction between *Tc*MRGBP and *Tc*BDF6**. Spot growth assay of MaV203 strains expressing a combination of DB, AD, AD-*Tc*BDF6, DB-*Tc*MRGBP, AD-*Tc*MRGx and GFP-*Tc*MRGx. The table on the left indicates the combinations tested in each lane. Positive and negative interaction controls were also included. Dilutions of $OD_{600}$=1.000, 0.100, 0.010 and 0.001 were spotted on solid SC medium or solid SC medium without histidine (-His) supplemented with 50mM 3-AT to titrate basal HIS3 expression. Plates were incubated at 30 °C for 6 days. In this case we used a complete set of experimental controls, in particular the GFP control, to test if the relative increase in *HIS3* expression was a non-specific engagement with GFP.

### Figure S7

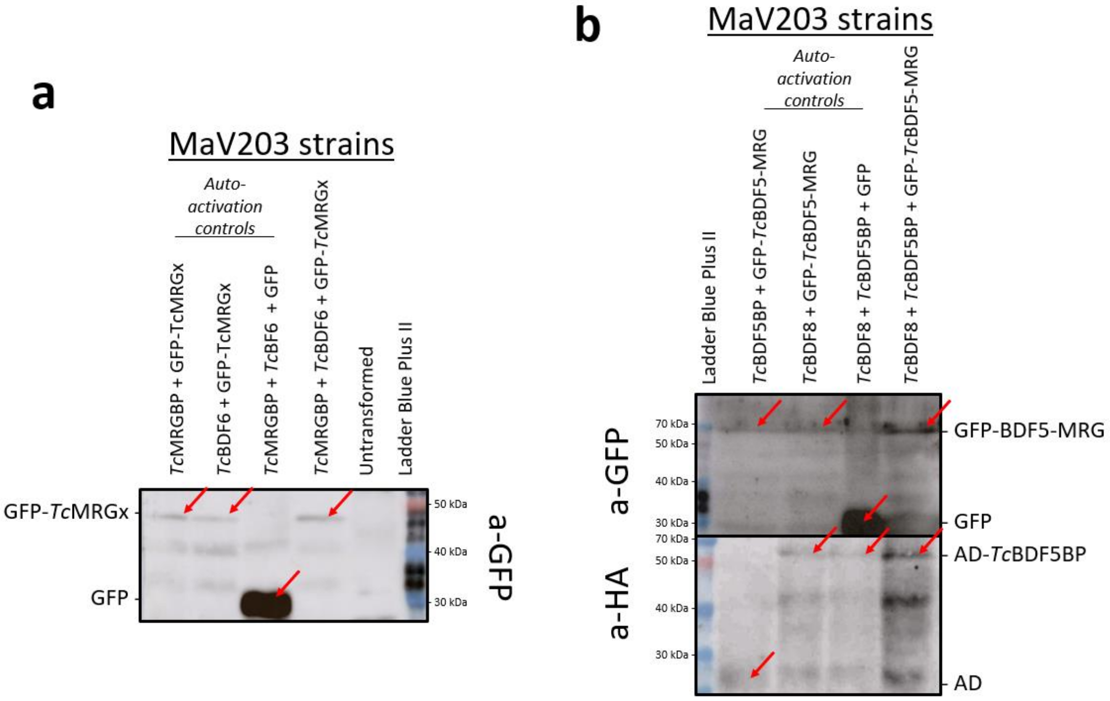


**Figure S7: Western blots showing the expression of HA and GFP-tagged proteins in the strains for Y3H experiments.** ***(a)*** Western blot showing the expected sizes of the third protein introduced to each strain for the *Tc*TINTIN Y3H experiment of **Figure S6**. **(b)** Expected sizes of GFP and HA-tagged proteins on yeast strains for the Y3H experiment of *Tc*BDF5-MRG heterotrimer.

### Figure S8

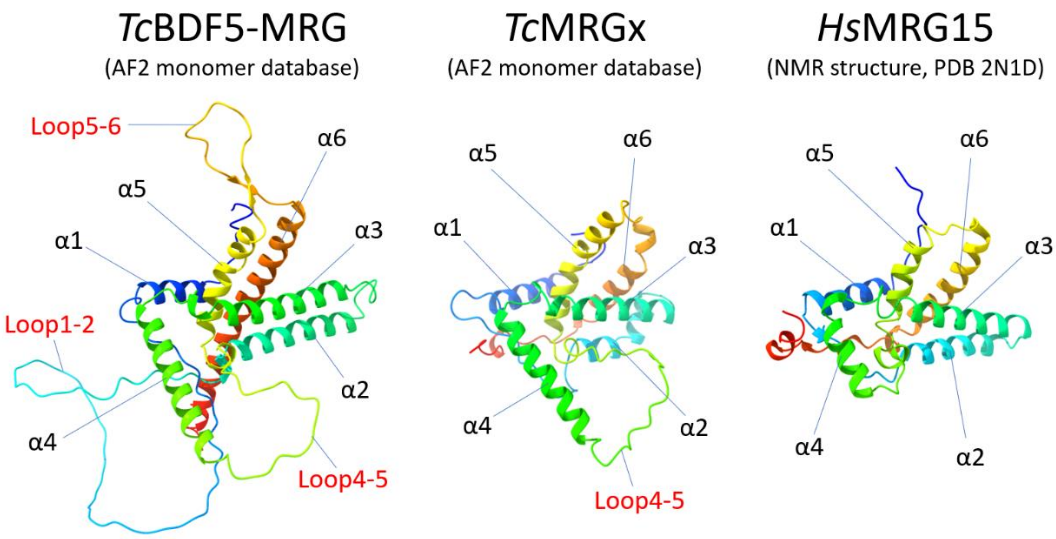


**Figure S8: Structure comparison between the MRG domain of *Tc*BDF5 and the MRG domain of *Hs*MRG15 and *Tc*MRGx**. The MRG of *Tc*BDF5 presents one mayor insertion of approximately 50 aminoacids between alpha helices 1 and 2 (Loop1-2), and two minor insertions, one of which is also present in *Tc*MRGx (Loop4-5), but not the other (Loop5-6). Structures were aligned to *Tc*MRGx using matchmaker algorithm in ChimeraX.

### Figure S9

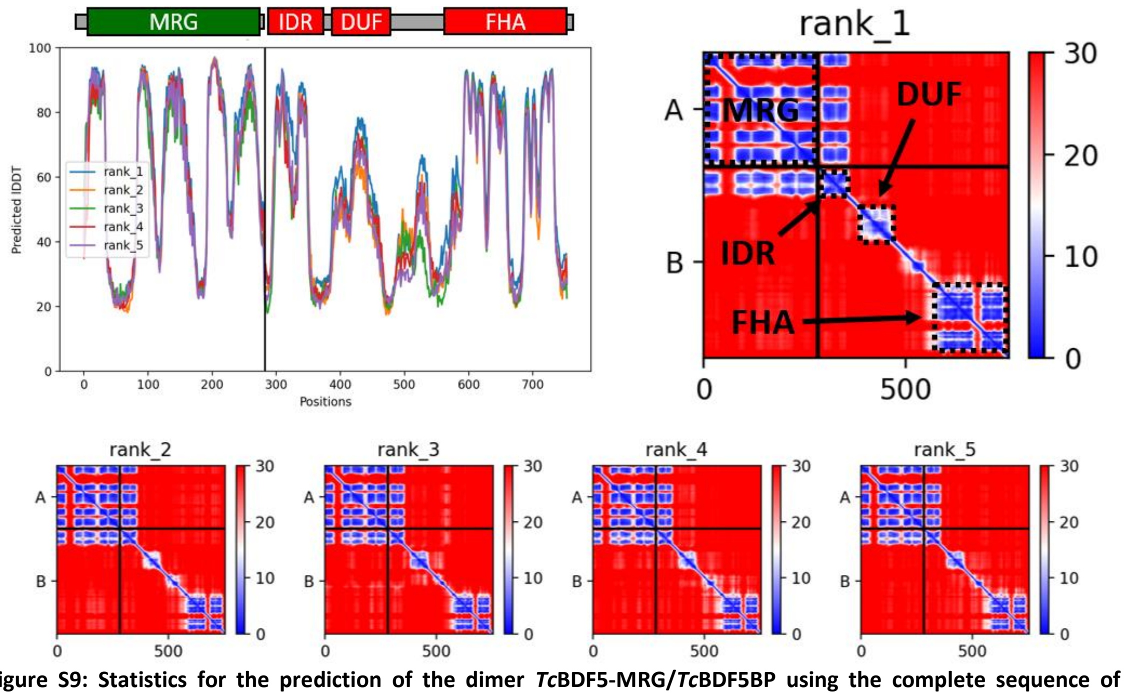


**Figure S9: Statistics for the prediction of the dimer *Tc*BDF5-MRG/*Tc*BDF5BP using the complete sequence of *Tc*BDF5BP**. The PAE graph shows that only the IDR of *Tc*BDF5BP is involved in the interaction.

### Figure S10

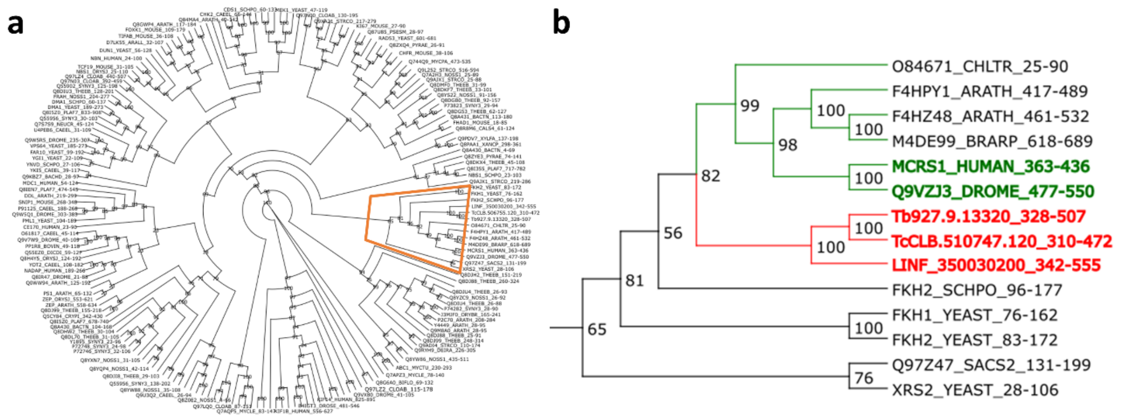


**Figure S10: Phylogenetic tree of FHA domains. (a)** Complete phylogenetic tree of the FHA domains sequences from the Pfam entry PF00498, adding the sequences of FHA domains of BDF5BPs from *T. brucei*, *T. cruzi* and *L. infantum*. The orange rectangle encloses the subtree from figure b. **(b)** IDs and branches from trypanosomatids BDF5BPs are shown in red. Green branches correspond to FHA proteins that cluster with trypanosomatids BDF5BP. The bold green IDs are characterized proteins similar in size and architecture to trypanosomatids BDF5BP. The IDs are followed by an underscore and the limits used for each protein to generate the tree.

### Figure S11

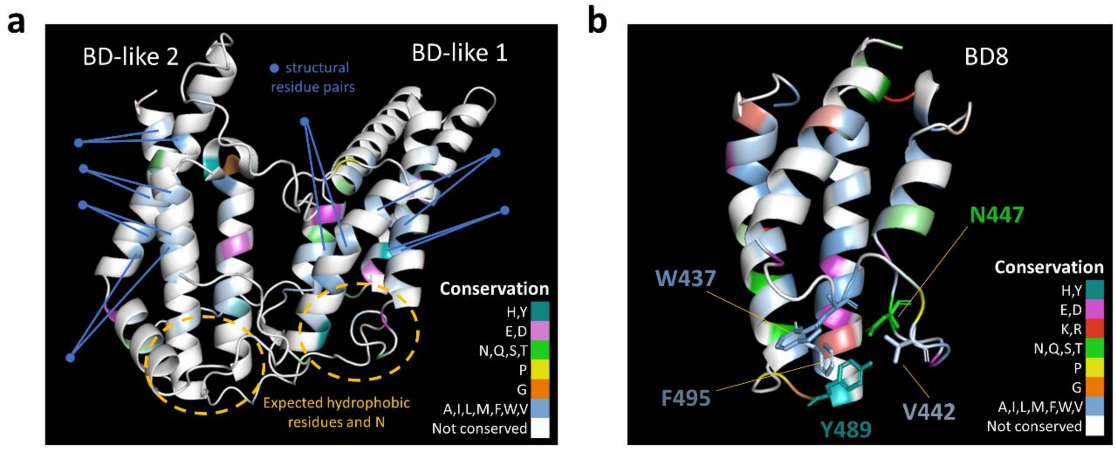


**Figure S11: Conserved residues of TcBDF8 domains.** BD and BD-like structures present in *Tc*BDF8. Conserved residues are colored by conservation and by physicochemical properties. Not conserved residues are in white. Brighter colors refer to a higher degree of conservation. (**a**) Bromodomain-like structures from the DUF domain. The pairs of blue lines coming from the blue dots shows the pairs of conserved hydrophobic residues that may be involved in keeping the fold of the domain. The yellow circles points to the region in which must be located the conserver residues involved in acetyl-lysine recognition. (**b**) C-terminal bromodomain (BD8). The region involved in acetyl-lysine recognition has many conserved residues that could be involved in this function. The position 489 is expected to have a conserved asparagine, but instead it has a conserved tyrosine.

**Figure S12**

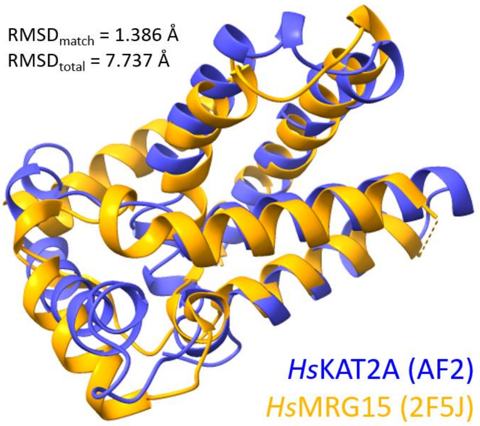


**Figure S12: Structural similarity between residues 227 to 370 of human KAT2A and the MRG domain of human MRG15.** The residues 227-370 of KAT2A predicted structure (AF2) were structurally aligned to the crystal structure of human MRG15 (PDB: 2F5J). An RMSD of 1.386 Å was obtained between 26 pruned atom pairs and of 7.737 Å for the entire structures.

### Supporting table

**Table S1: Primers used in the study.**

| Primer | Gene name | Gene ID | Sequences |
|---|---|---|---|
| TcMRGx_BamHI-Fw | *Tc*MRGx | C4B63_22g316 | AACGGATCCATGGAGGGAGTTGACTGG |
| TcMRGx_XhoI-Rv | *Tc*MRGx | C4B63_22g316 | CCACTCGAGTCATTTCAAAGGAATATACGC |
| TcMRGx-R138A-Fw | *Tc*MRGx | C4B63_22g316 | GTGGAGTATCTTCTTGCCTTTTTTATGGCTTTGC |
| TcMRGx-R138A-Rv | *Tc*MRGx | C4B63_22g316 | GCAAAGCCATAAAAAAGGCAAGAAGATACTCCAC |
| TcMRGBP_SalI-Fw | *Tc*MRGBP | C4B63_55g378c | AAAGTCGACAATGCGATTGGCTACAAGTGG |
| TcMRGBP_XbaI-Rv | *Tc*MRGBP | C4B63_55g378c | AAATCTAGATTATCACTTTTCCTCACCCAGAAGC |
| TcBDF6-HA_BamHI-Fw | *Tc*BDF6 | BCY84_19389 | AAAGGATCCATGTATCCGTATGATGTGCCGGATTATG CTCGGCGGGAAGATTTACTGC |
| TcBDF6_EcoRV-Rv | *Tc*BDF6 | BCY84_19389 | AAAGATATCTCATGCACCACGCAAATGC |
| TcBDF5-MRG_BamHI-Fw | *Tc*BDF5 | C4B63_1g51 | TTGGATCCCCTCCGCTACGGGCTCC |
| TcBDF5-MRG_XhoI-Rv | *Tc*BDF5 | C4B63_1g51 | AAACTCGAGCTACGATATTTCTTTTCCAATTTCCG |
| TcBDF5-R623A-Fw | *Tc*BDF5 | C4B63_1g51 | CTTGTGTTATCTTGTTGCTTTTTTGCAGCACCTTC |
| TcBDF5-R623A-Rv | *Tc*BDF5 | C4B63_1g51 | GAAGGTGCTGCAAAAAAGCAACAAGATAACACAAG |
| TcBDF5BP_SalI-Fw | *Tc*BDF5BP | C4B63_2g92 | AAAGTCGACAATGGATGGGTCGCGGTTTGG |
| TcBDF5BP_XbaI-Rv | *Tc*BDF5BP | C4B63_2g92 | AAATCTAGATTATTAATCCTCCATCAACACCTCGG |
| TcBDF5BP-IDR_XbaI-Rv | *Tc*BDF5BP | C4B63_2g92 | AAATCTAGATTATGTCGTCATGGGTGCTGCAG |
| TcBDF5BP-NoIDR_XbaI-Fw | *Tc*BDF5BP | C4B63_2g92 | AAAGTCGACAATGCTGGCCTCATTGTTGGCCG |
| TcBDF8_SalI-Fw | *Tc*BDF8 | C4B63_7g80 | AAAGTCGACAATGGATTTGATTTTTGCTGAGGGAG |
| TcBDF8_XbaI_Rv | *Tc*BDF8 | C4B63_7g80 | AAATCTAGATTATCACAACAGGCCCACCTCG |
| TcBDF8-Peptide_XbaI-Rv | *Tc*BDF8 | C4B63_7g80 | AAATCTAGATTATGGCAGGACTCGTGCCGC |
| TcBDF8-DUF_SalI_Fw | *Tc*BDF8 | C4B63_7g80 | AAAGTCGACAATGGCGGCACGAGTCCTGCCAG |
| TcBDF8-DUF_XbaI_Rv | *Tc*BDF8 | C4B63_7g80 | AAATCTAGATTAAGGAAGTGGAGTCGGCGTC |
| TcBDF8_BD8_SalI-Fw | *Tc*BDF8 | C4B63_7g80 | AAAGTCGACAATGTCCTGCACTGATACACAGC |

### Molecular dynamics simulations

This section includes several figures and analysis about the performed molecular dynamics that are not discussed in the main text.

In order to study the structural variation of *Tc*MRGBP on different states (monomer and heterodimer), we performed molecular dynamics simulations. According to our results,

*Tc*MRGBP has a high residue fluctuation when it is separated from *Tc*MRGx, especially in the IDR region (residues 101-125 in gray).

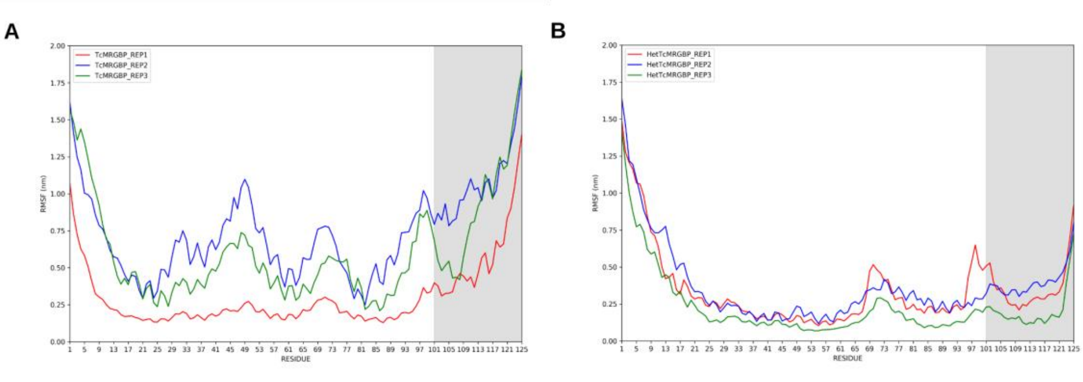


**RMSF plots**. RMSF values calculated by residue for each conformation from MD simulation replicas. It is observed that the IDR region (101-125 residues) has more fluctuation in TcMRGBP monomer (A) structure than TcMRGx/TcMRGBP dimer (B).

We also observed that spite of the fact that both *Tc*MRGBP states have shown high structural variation compared to model obtained with AlphaFold program, the averages of RMSD values for each replica of *Tc*MRGBP in the monomeric condition were higher than those values obtained on *Tc*MRGBP in complex with *Tc*MRGx:

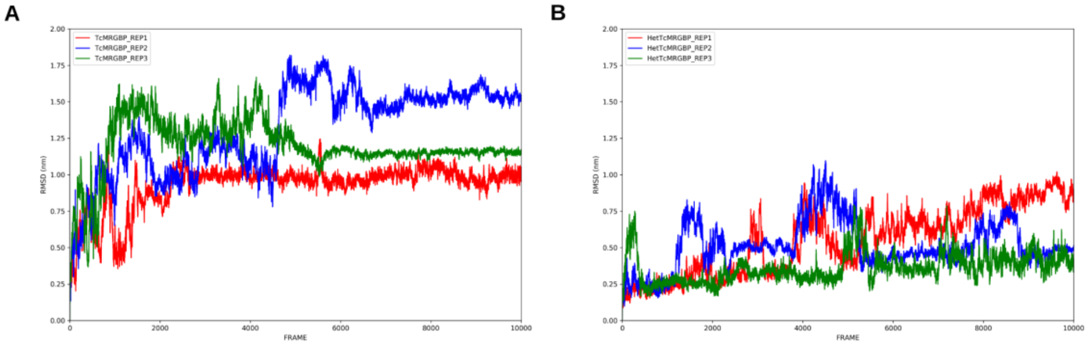


**RMSD plots**. RMSD values calculated for each conformation from MD simulation replicas. Despite the fact that both *Tc*MRGBP in monomer (A) and *Tc*MRGBP in dimer (B) have high fluctuations in structure. *Tc*MRGBP in the monomer presents higher RMSD averages than *Tc*MRGBP in the heterodimer.

Radius of gyration (RoG) and total surface area solvent accessible (SASA) changes also corroborate that *Tc*MRGx is important to *Tc*MRGBP structure stabilization:

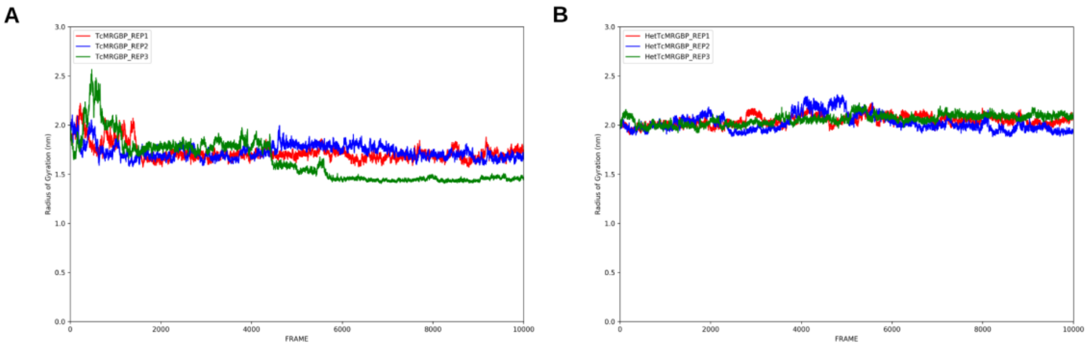


**RoG plots**. RoG values calculated for each conformation from MD simulation replicas.
It is observed that TcMRGBP in monomer (A) has a subtle and small variation of the RoG averages when compared to TcMRGBP in dimer (B). Differences in TcMRGBP values in the monomer may indicate structural changes in relation to the protein's center of mass.

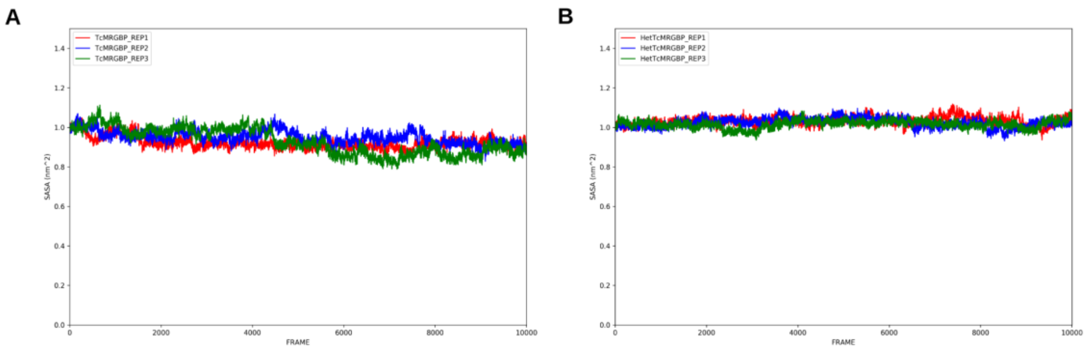


**SASA plots**. SASA values calculated for each conformation from MD simulation replicas. It is observed that TcMRGBP in monomer (A) has a subtle and small variation of the SASA averages when compared to TcMRGBP in dimer (B). These values may indicate that TcMRGBP in monomer suffers changes on its folding to reduce the exposition on solvent and consequently compacts the structure when compared to TcMRGBP in dimer.

In the RoG plot, we may see that RoG values obtained for *Tc*MRGBP in the monomeric condition have a subtle and small variation on the replica averages compared to *Tc*MRGBP in the heterodimeric condition. Despite the subtle differences, ANOVA test indicated a significant variation in the replica averages between two *Tc*MRGBP structural states. These differences in RoG values observed between monomeric and heterodimeric states may indicate important structural changes in relation to the protein's center of mass. Considering SASA values (SASA plot), *Tc*MRGBP in monomer replicas also showed a subtle, but significant variation in surface area compared to target structure in heterodimer. The reduction on SASA values observed in *Tc*MRGBP in monomer state replicas is a consequence of a fold change which induces a reduction on exposed residues to solvent and protein structure compaction. This can be seen in the WHAM plots:

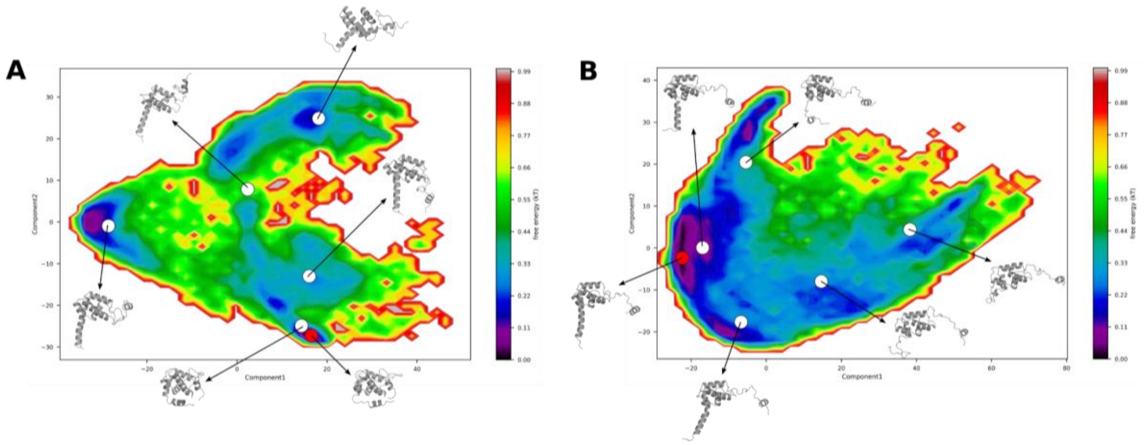


**WHAM plots**. Free-energy map calculated using RMSD as reaction coordinate. In (A) we present the free-energy map of *Tc*MRGBP in the monomer condition with medoid structures obtained with agglomerative ward algorithm. In (B) we present the free-energy map of *Tc*MRGBP in the heterodimeric condition and medoid structures obtained with agglomerative ward algorithm. Structures on the red circle represent the structure on the minimum free-energy state.

Our results suggest that *Tc*MRGx is necessary for *Tc*MRGBP structure stabilization and also suggest the existence of a conformational change on *Tc*MRGBP between both states. These effects combined can be ultimately responsible for the discrepancy between predicted interactions using AF2-multimer models and the experimental evidence obtained by Y2H for *Tc*MRGBP and *Tc*BDF6.
